# m^6^A mRNA Methylation in Brown Adipose Tissue Regulates Systemic Insulin Sensitivity via an Inter-Organ Prostaglandin Signaling Axis

**DOI:** 10.1101/2023.05.26.542169

**Authors:** Ling Xiao, Dario F. De Jesus, Cheng-Wei Ju, Jiang-Bo Wei, Jiang Hu, Ava DiStefano-Forti, Tadataka Tsuji, Cheryl Cero, Ville Männistö, Suvi M. Manninen, Siying Wei, Oluwaseun Ijaduola, Matthias Blüher, Aaron M. Cypess, Jussi Pihlajamäki, Yu-Hua Tseng, Chuan He, Rohit N. Kulkarni

## Abstract

Brown adipose tissue (BAT) has the capacity to regulate systemic metabolism through the secretion of signaling lipids. N6-methyladenosine (m^6^A) is the most prevalent and abundant post-transcriptional mRNA modification and has been reported to regulate BAT adipogenesis and energy expenditure. In this study, we demonstrate that the absence of m^6^A methyltransferase-like 14 (METTL14), modifies the BAT secretome to initiate inter-organ communication to improve systemic insulin sensitivity. Importantly, these phenotypes are independent of UCP1-mediated energy expenditure and thermogenesis. Using lipidomics, we identified prostaglandin E2 (PGE2) and prostaglandin F2a (PGF2a) as M14^KO^-BAT-secreted insulin sensitizers. Notably, circulatory PGE2 and PGF2a levels are inversely correlated with insulin sensitivity in humans. Furthermore, *in vivo* administration of PGE2 and PGF2a in high-fat diet-induced insulin-resistant obese mice recapitulates the phenotypes of METTL14 deficient animals. PGE2 or PGF2a improves insulin signaling by suppressing the expression of specific AKT phosphatases. Mechanistically, METTL14-mediated m^6^A installation promotes decay of transcripts encoding prostaglandin synthases and their regulators in human and mouse brown adipocytes in a YTHDF2/3-dependent manner. Taken together, these findings reveal a novel biological mechanism through which m^6^A-dependent regulation of BAT secretome regulates systemic insulin sensitivity in mice and humans.

**Highlights:**

- Mettl14^KO^-BAT improves systemic insulin sensitivity via inter-organ communication;
- PGE2 and PGF2a are BAT-secreted insulin sensitizers and browning inducers;
- PGE2 and PGF2a sensitize insulin responses through PGE2-EP-pAKT and PGF2a-FP-AKT axis;
- METTL14-mediated m^6^A installation selectively destabilizes prostaglandin synthases and their regulator transcripts;
- Targeting METTL14 in BAT has therapeutic potential to enhance systemic insulin sensitivity

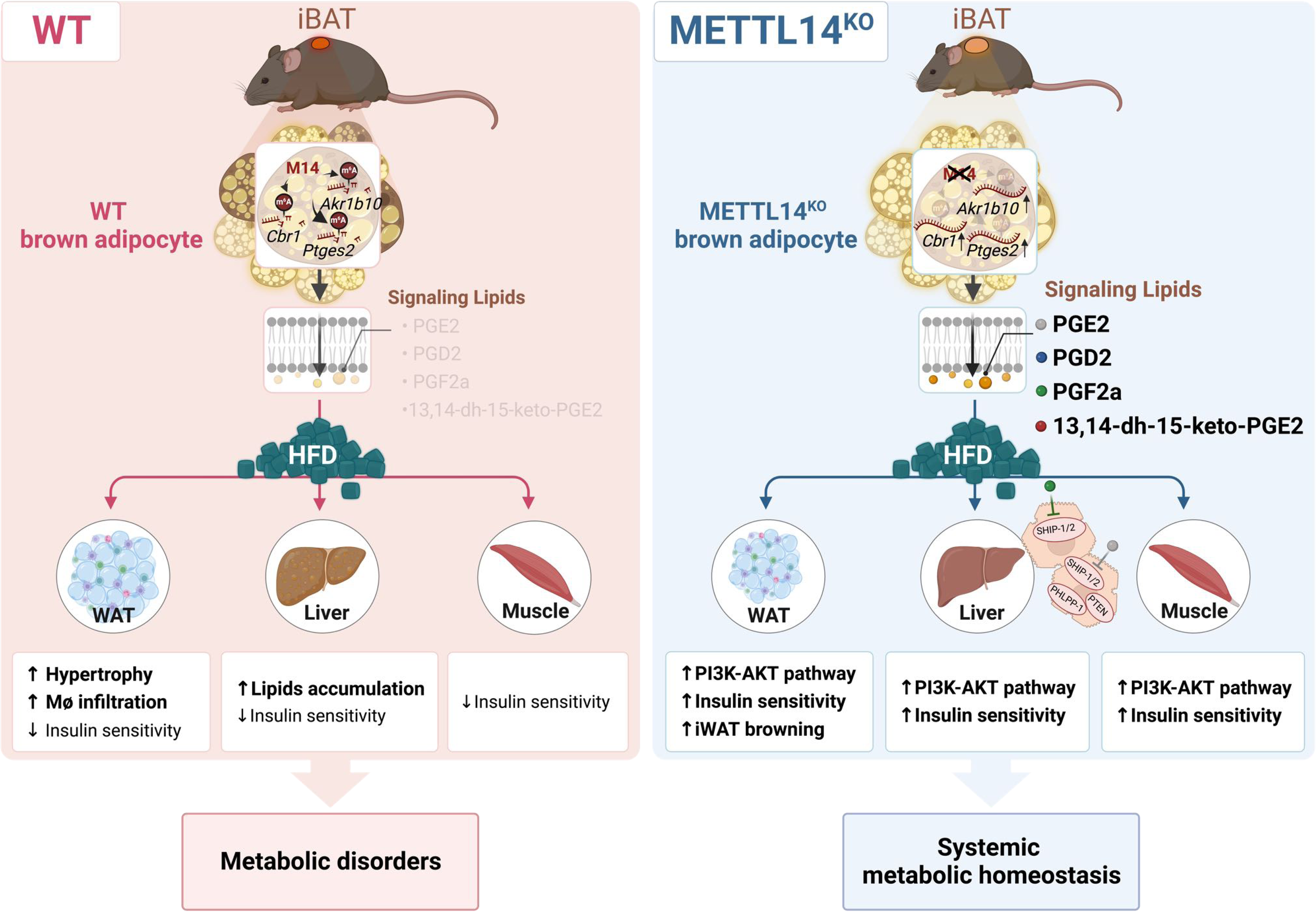

## Introduction

Brown adipose tissue (BAT) has emerged as a potential target for the treatment and prevention of human obesity and related metabolic disorders secondary to its thermogenic function mediated by uncoupling protein 1 (UCP1). In humans, BAT-mediated energy expenditure under cold exposure contributes to body fat reduction (Ouellet et al. 2012; Yoneshiro et al. 2013). BAT has also been reported to promote cardiometabolic health, especially in overweight or obese individuals (Becher et al. 2021). Although BAT has been traditionally recognized for its thermogenic capacity upon cold stimulation, it can also provide metabolic benefits by its ability to assist in utilization of nutrients (e.g., glucose, fatty acids, branched chain amino acids) during energy dissipation, and via actions of secreted factors (Shamsi, Wang, and Tseng 2021).

Notably, BAT has been reported to exert positive effects on systemic metabolism via secreted factors that include peptides, proteins, lipids, or microRNAs (Shamsi, Wang, and Tseng 2021; Cypess 2022). In particular, a class of lipids known as ‘brown lipokines’ has attracted attention in recent years (Shamsi, Wang, and Tseng 2021; Cypess 2022). The term ‘lipokine’ was first introduced to define a type of adipose tissue-secreted lipid hormone linking to systemic metabolic processes, including the regulation of insulin sensitivity, glucose tolerance, and inflammation (Cao et al. 2008). Several adipokines have been identified, for example, 12,13-dihydroxy-octadecaenoic acid (12,13-diHOME), a cold- and exercise-induced BAT-derived lipokine, has been shown to increase fatty acid transport into BAT, leading to decreased circulating triglycerides (Lynes et al. 2017), and to promote fatty acid uptake in skeletal muscle (Stanford et al. 2018). Another cold-induced BAT-secreted signaling lipid, 12-hydroxy-eicosapentaenoic acid (12-HEPE), acts in an autocrine and endocrine fashion to promote glucose uptake into BAT and skeletal muscle (Leiria et al. 2019), while BAT-secreted Maresin 2 acts as an anti-inflammatory lipid to mediate BAT-liver crosstalk (Sugimoto et al. 2022). Nonetheless, virtually all signaling lipids secreted by BAT have been identified via manipulations that include cold exposure, exercise, or β-adrenergic agonists, with virtually no reports interrogating the significance of the epitranscriptome on the BAT secretome.

N6-methyladenosine (m^6^A) is the most prevalent, abundant and conserved internal post-transcriptional modification in eukaryotic RNAs (X. Wang, Lu, et al. 2014). The modification is primarily orchestrated by the writer complex, which consists of methyltransferase-like 14 (METTL14), methyltransferase-like 3 (METTL3) and Wilms’ tumor 1-associated protein (WTAP) (Liu et al. 2014). Removal of m^6^A is performed by demethylases such as fat mass and obesity-associated protein (FTO) or alkB homologue 5 (ALKBH5). The effects of the m^6^A modification on mRNA metabolism depend largely on the recognition by different m^6^A reader proteins to allow regulation of mRNA stability, translation, splicing, and/or export. For instance, recognition of m^6^A by both YTH N(6)-methyladenosine RNA binding protein 2 (YTHDF2) or YTH N(6)-methyladenosine RNA binding protein 3 (YTHDF3) controls mRNA decay of the targeted mRNAs (Shi et al. 2017; X. Wang, Lu, et al. 2014).

While recent studies have implicated m^6^A modification in diverse physiological processes including the regulation of human β-cell biology (De Jesus et al. 2019) and in brown/beige adipogenesis and/or thermogenesis (Y. Wang et al. 2020; Claussnitzer et al. 2015; Yan et al. 2023), its significance in modifying the BAT secretome to regulate systemic insulin sensitivity remains unexplored.

In the present study, we propose that METTL14 regulates BAT secretory function by a unique mechanism to regulate systemic insulin sensitivity. BAT-specific METTL14 knockout (M14^KO^) mice displayed improved insulin sensitivity and glucose tolerance independent of body weight, sex or canonical BAT thermogenesis. Untargeted lipidomics profiling of mouse BAT and human brown adipocytes (hBAT) identified upregulation of PGE2, PGD2, PGF2a, and 13,14-dh-15-keto-PGE2 secondary to METTL14 deficiency. Mechanistically, PGE2 and PGF2a improve systemic insulin sensitivity via suppressing AKT phosphatases in key peripheral metabolic tissues. Conversely, METTL14 ablation in hBAT prevents the decay of specific transcripts encoding prostaglandin synthases to enable an increase in their protein levels.

Taken together, these results argue for a novel mechanism coupling m^6^A methylation with the BAT-secretome, independent of thermogenesis, and suggest METTL14 as a potential therapeutic target for improving systemic insulin resistance.

## Results

### *Mettl14* Deficiency in BAT Improves Systemic Insulin Sensitivity and Glucose Tolerance Independent of UCP1-mediated Thermogenesis or Energy Expenditure in Mice

First, to determine whether expression of m^6^A writers is altered in obesity and insulin sensitivity, we performed qRT-PCR analyses in human brown adipose tissue biopsies obtained from lean (BMI<30) or obese (BMI ≥30) humans. Gene expression of *METTL14*, but not *METTL3* and *WTAP*, was dramatically upregulated in BAT from obese compared to lean individuals (Figure 1A). To further investigate the involvement of BAT m^6^A in systemic insulin sensitivity, we examined mouse models of systemic insulin resistance. We analyzed the expression levels of m^6^A writers *Mettl14*, *Mettl3*, and *Wtap* in the interscapular BAT (iBAT) samples of a genetic hyperphagic mouse model of insulin resistance (Coleman 1978), namely, the leptin receptor-deficient *db/db* mouse. *db/db* mice exhibited significantly higher Mettl14 at both mRNA (Figure 1B) and protein (Figures 1C and S1A) levels compared to controls. Next, to distinguish insulin sensitivity from obesity, we took advantage of the liver-specific insulin receptor knockout (LIRKO) mouse model, which exhibits severe insulin resistance in the absence of obesity (Michael et al. 2000). Again, *Mettl14* gene and protein expression levels in iBAT were upregulated in the LIRKOs compared to littermate controls (Figures 1D, 1E, and S1B). These findings suggest a potential role for METTL14 in modulating BAT to regulate systemic insulin sensitivity and metabolic homeostasis.

**Figure 1.**
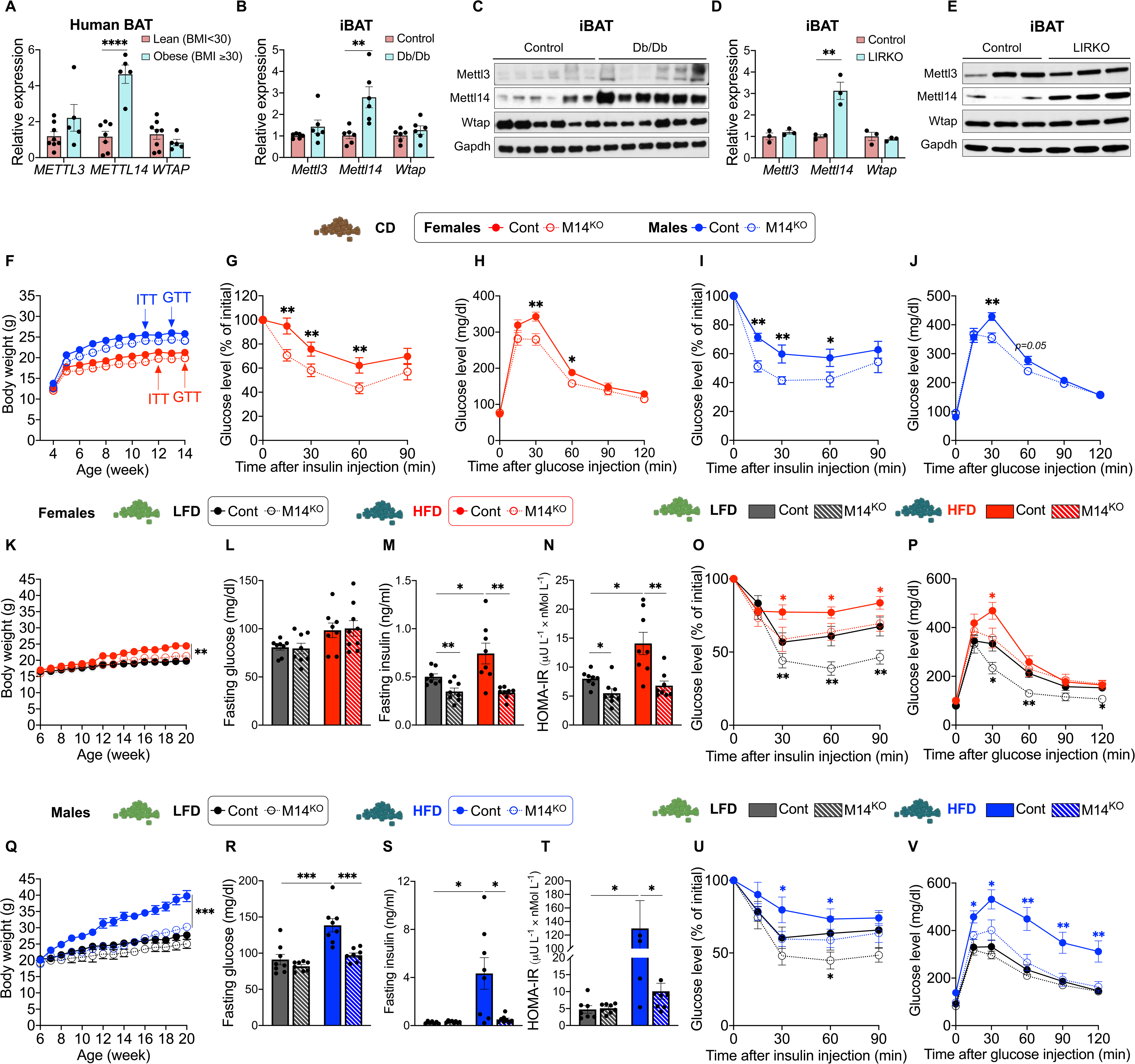
Ablation of Mettl14 in BAT Improves Systemic Insulin Sensitivity and Glucose Tolerance in Mice. (A) qRT-PCR of m^6^A writer’s genes *METTL3*, *METTL14*, and *WTAP* in human brown adipose tissues (n=8 for lean; n=5 for obese). (B) qRT-PCR of *Mettl3*, *Mettl14*, and *Wtap* in interscapular brown adipose tissues of control and *db/db* mice (n=6/group). (C) Western blots of Mettl3, Mettl14, Wtap, and Gapdh in interscapular brown adipose tissues of control and *db/db* mice (n=6/group). (D) qRT-PCR of *Mettl3*, *Mettl14*, and *Wtap* in interscapular brown adipose tissues of control and LIRKO mice (n=3/group). (E) Western blots of Mettl14 and Gapdh in interscapular brown adipose tissues of control and LIRKO mice (n=3/group). (F) Body weight trajectories of CD-fed mice (females: n=12 for control; n=9 for M14^KO^ groups; males: n=10 each for control and M14^KO^ groups). (G and H) Insulin (G) and glucose tolerance tests (H) of CD-fed females (n=12 for control, and n=9 for M14^KO^ groups). (I and J) Insulin (I) and glucose tolerance tests (J) of CD-fed males (n=10 each for control, and M14^KO^ groups). (K and G) Body weight trajectories of LFD- or HFD-fed control and Mettl14^KO^ females (K) and males (G) (females; n=8 for control, and n=5 for M14^KO^ groups on LFD; females; n=8 for control and n=6 for Mettl14^KO^ groups on HFD; and males: n=8 each for control and M14^KO^ groups on LFD or HFD). (L and R) Fasting glucose levels of control and M14^KO^ females (L) and males (R) (n=7-8 for each group). (M and S) Fasting insulin levels in the serum of control and M14^KO^ females (M) and males (S) (n=7-8 for each group). (N and T) HOMA-IR of LFD- or HFD-fed control and M14^KO^ females (N) and males (T) (n=7-8 for each group). (O and U) Insulin tolerance tests of LFD- or HFD-fed control and M14^KO^ females (O) and males (U) (n=7-8 for each group). (P and V) Glucose tolerance tests of LFD- or HFD-fed control and M14^KO^ females (P) and males (V) (n=7-8 for each group). All samples in each panel are biologically independent. Data are expressed as means ± SEM. *p < 0.05, **p < 0.01, ***p < 0.001 by Two-way ANOVA (F-J, K, O-Q, U and V) and Two-tailed unpaired t-test (A, B, L-N, and R-T). Also see Figure S1.

To directly interrogate the role of Mettl14, we generated BAT-specific Mettl14 knockout mice by crossing *Mettl14*-floxed (M14^fl/fl^) animals (De Jesus et al. 2019) with *Ucp1*-cre mice (Kong et al. 2014). We confirmed that Mettl14 was specifically depleted in iBAT and its protein abundance and gene expression were not significantly altered in other metabolic tissues including iWAT, eWAT, liver or muscle of *M14*^fl/fl^-*Ucp1*-cre (referred as M14^KO^) mice compared to controls (M14^fl/fl^) (Figure S1C-S1E). The KO and control mice were born in a normal Mendelian ratio and did not show developmental defects (data not shown).

To explore the role of Mettl14 under physiological conditions, we first placed both female and male mice on a regular chow diet (CD). Ablation of Mettl14 did not significantly change the body weight of male and female mice fed with CD (Figure 1F). However, both sexes of M14^KO^ mice showed improved insulin sensitivity and glucose tolerance (Figures 1G-J).

To further examine the ability of M14^KO^ mice to handle insulin resistance we fed mice with HFD (60% fat) and compared with mice fed low fat diet (LFD) (10% fat) as a controls. M14^KO^ mice on HFD were smaller in body size (Figure S1F) and exhibited increased iBAT but decreased iWAT, eWAT and liver mass (Figure S1G). In line with these findings, hematoxylin and eosin (H&E) staining showed larger brown adipocytes (Figure S1H) while the size of adipocytes in the inguinal (Figure S1I) and epididymal white fat were smaller (Figure S1J) in LFD-fed M14^KO^ mice. Interestingly, although HFD increased the brown adipocyte size in controls, as expected, it did not significantly alter the size of brown adipocytes in the M14^KO^ mice (Figure S1H), suggesting an increase in their adipocyte numbers in the mutants. The size of adipocytes in the inguinal (Figure S1I) and epididymal white fat (Figure S1J) from HFD-M14^KO^ mice were visually smaller compared to controls. Surprisingly, immunohistochemical (IHC) staining displayed lower UCP1 abundance in M14^KO^ iBAT (Figure S1H). The presence of UCP1-positive cells in the iWAT of LFD-fed M14^KO^ mice (Figure S1I), suggested browning of white adipose tissue. In contrast to controls, M14^KO^ mice were protected from HFD-induced macrophage infiltration as indicated by F8/40 IHC staining in eWAT (Figure S1J). Furthermore, M14^KO^ mice were protected from HFD-induced hepatic steatosis (Figure S1K). The morphological changes in iWAT and eWAT were accompanied by smaller adipocyte diameters in male M14^KO^ mice on HFD compared to controls (Figures S1L and S1M).

As expected, control female mice fed a HFD presented increased body weights, elevated fasting insulin levels, and insulin resistance as measured by the homeostatic model assessment of insulin resistance (HOMA-IR) compared to control female mice fed a LFD (Figures 1K-P). In contrast, female M14^KO^ mice were protected against HFD-induced insulin resistance and presented lower fasting insulin levels and improved HOMA-IR compared to controls on HFD (Figures 1K-P). Thus, female M14^KO^ mice presented improved insulin sensitivity and glucose tolerance profiles independent of the diet compared to controls (Figures 1R and S). Similar changes were observed in males (Figures 1Q-V).

Serum adiponectin levels were comparable between M14^KO^ mice and littermate control counterparts in both females (Figure S1N) and males (Figure S1O). Circulating leptin levels were lower in the M14^KO^ mice on a HFD in females (Figure S1N) and males (Figure S1O). Overall, these data suggest that Mettl14 ablation in BAT improves insulin sensitivity and glucose tolerance independent of body weight or diet.

Finally, we examined the contribution of UCP1-dependent thermogenesis in the observed phenotypes of M14^KO^ mice. The reduced abundance of UCP1 suggested by IHC in the M14^KO^-iBAT (Figure S1H) was confirmed by Western blot analyses in the iBAT from M14^KO^ mice on both diets compared to corresponding controls (Figures S1P). This prompted us to hypothesize that the improvement in insulin sensitivity and glucose tolerance in M14^KO^ mice occurs independent of the canonical BAT function, namely UCP1-mediated thermogenesis and energy expenditure. While we observed improved cold tolerance in M14^KO^ males (Figure S1Q), we did not detect differences in energy expenditure (Figure S1R), oxygen consumption (Figure S1S), or carbon dioxide production (Figure S1T), as measured by indirect calorimetry, between the control and M14^KO^ groups.

Taken together, these results suggest METTL14 acts to limit BAT-mediated whole-body metabolism in normal physiology and this occurs independent of a reduction in body weight as well as intrinsic thermogenetic capacity of the BAT.

### M14^KO^-BAT-secreted Factors Enhance Insulin Sensitivity in Peripheral Metabolic Tissues in Mice and Primary Metabolic Cells in Humans

The improved insulin sensitivity following ablation of Mettl14 in iBAT that was independent of gender, body weight or diet promoted us to examine the tissue-specific insulin sensitizing effects using *in vivo* insulin stimulation experiments (Figure 2A). To this end, we performed *in vivo vena cava* insulin infusion in the various groups followed by harvesting of metabolic tissues post-insulin stimulation (Figure 2A). M14^KO^ mice displayed a dramatically higher hepatic serine 473 phosphorylation of protein kinase B (pAkt_S473_) compared to controls independent of diet or insulin-stimulated phosphorylation of insulin or Igf1 receptors (Irβ/Igf1rβ) (Figures 2B and 2C).

**Figure 2.**
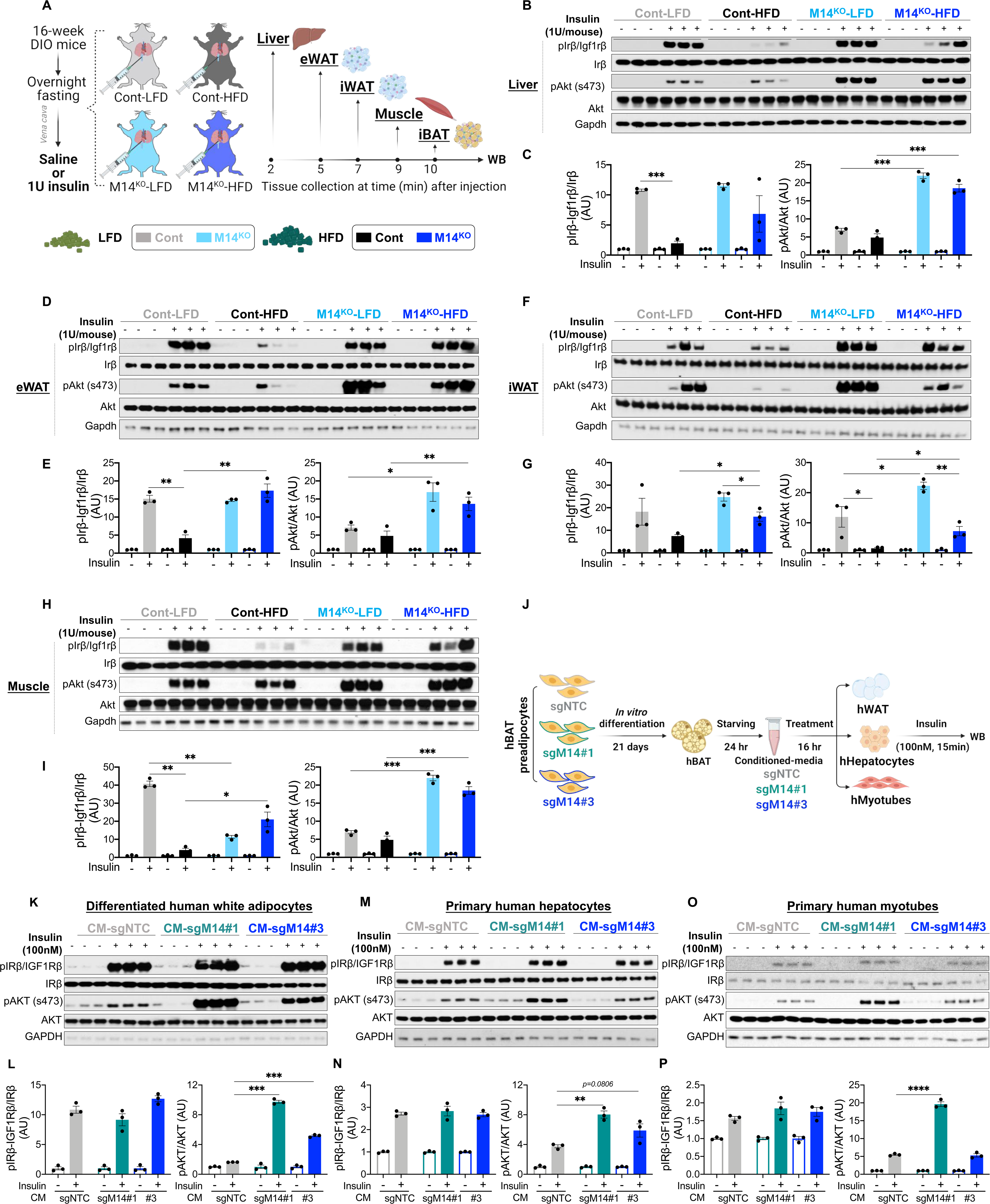
Ablation of Mettl14 in BAT Enhances Insulin Sensitivity of Mouse Peripheral Metabolic Tissues and Human Metabolic Cells by Secretory Factors. (A) Schematic of metabolic tissues collection at indicated timepoints after *vena cava* saline/insulin injection in LFD- or HFD-fed control and M14^KO^ male mice (n=6/group). (B-I) The insulin-stimulated phosphorylation of Ir/Igf1rβ and Akt_S473_ in liver (B and C), eWAT (D and E), iWAT (F and G), and muscle (H and I) after injection of 1U insulin into the *vena cava*. (J) Experimental scheme of *in vitro* co-culture experiments. Differentiated human white adipocytes, human primary hepatocytes, and differentiated human myotubes were treated with differentiated human brown adipocytes conditioned-media for 16hr and stimulated with 100 nM insulin for 15 min. (K-P) The insulin-stimulated phosphorylation of IR/IGF1Rβ and AKT_S473_ in hBAT conditioned media treated human white adipocytes (K and L), human primary hepatocytes (M and N), and differentiated human myotubes (O and P) (n=3 independent experiments). All samples in each panel are biologically independent. Data are expressed as means ± SEM. *p < 0.05, **p < 0.01, ***p < 0.001 by Two-tailed unpaired t-test (B, E, G, I, L, N, P). CM, conditioned media. Also see Figure S2.

Additionally, when fed a LFD, M14^KO^ mice exhibited elevated pAkt_S473_ in both their eWAT (Figures 2D and 2E) and iWAT (Figures 2F and 2G) compared to controls. Furthermore, the typical decrease in pIrβ/Igf1rβ and pAkt_S473_ that occurs in response to HFD, and that was evident in controls, was not detected in eWAT (Figures 2D and 2E) or iWAT (Figures 2F and 2G) from M14^KO^ mice.

While M14^KO^ mice fed with LFD showed reduced pIrβ/Igf1rβ in muscle, they demonstrated increased pAkt_S473_ activation compared to control mice on the same diet (Figures 2H and 2I). When challenged with HFD, M14^KO^ mice exhibited higher levels of both pIrβ/Igf1rβ and pAkt_S473_ in their muscle compared to control mice (Figures 2H and 2I).

Of note, no significant difference in pIrβ/Igf1rβ and pAkt levels were observed in iBAT between control and M14^KO^ mice on LFD (Figures S2A and S2B). While an increase in pIrβ/Igf1rβ was observed in M14^KO^-iBAT compared to control-iBAT in response to HFD, there were no differences in pAkt_S473_ levels between control- and M14^KO^-iBAT (Figures S2A and S2B). These findings suggest that the lack of Mettl14 in iBAT leads to increased insulin-stimulated responses primarily in the peripheral metabolic tissues, underscoring inter-organ communication in the M14^KO^ mouse model.

Our data thus far suggested the presence of secretory factor(s) generated by M14^KO^-BAT can enhance insulin sensitivity in distant metabolic organs, to improve systemic insulin sensitivity. To test this further, we generated three stable M14^KO^ human brown pre-adipocytes (sgM14#1-, sgM14#2-, and sgM14#3-hBAT) by using single guide RNA particles; single guide non-targeting control (sgNTC) RNA transfected hBAT cells were also generated as controls (Figure 2J). We verified METTL14 knockout in the differentiated hBAT cells *in vitro* by Western blot (Figure S2C). METTL14 depletion in hBAT cells resulted in an improved differentiation capacity as determined by Oil Red O staining (Figure S2D).

Next, we explored the relevance of METTL14 in regulating secretory function in hBAT cells. To mimic the *in vivo* setting, we performed co-culture experiments by treating differentiated human white adipocytes (hWAT), primary human hepatocytes (hHepatocytes), and differentiated primary human myotubes (hMyotubes) with serum-free conditioned-media from differentiated sgNTC- or sgM14-hBAT cells followed by analyses of the insulin-stimulated signaling of the recipient cells (Figure 2J). Consistent with our *in vivo* data, the conditioned media from sgM14-hBAT significantly increased pAKT_S473_ levels in differentiated hWAT cells in comparison to that from sgNTC-hBAT (Figures 2K and 2L). Consistently, conditioned media from sgM14-hBAT boosted pAKT_S473_ levels in hHepatocytes (Figures 2M and 2N) or hMyotubes (Figures 2O and 2P) in response to insulin when compared to cells treated with conditioned media from sgNTC-hBAT. Collectively, both *in vivo* and *in vitro* data strongly support our hypothesis that METTL14 ablation in BAT promotes the systemic release of brown adipocyte “factors” that promote intracellular insulin sensitization in distant metabolic tissues.

### Identification of Prostaglandin E2 (PGE2) and Prostaglandin F2 alpha (PGF2a) as the Major M14^KO^-BAT Secreted Insulin Sensitizers

We next sought to identify the putative factors secreted by brown adipocytes from M14^KO^mice. The ability of denatured hBAT-conditioned media to induce an improvement in pAKT_S473_ levels that was similar to the non-denatured conditioned media in hWAT cells (Figure S3A), allowed us to conclude that one or more factors released by the M14^KO^-brown adipocytes were likely signaling lipids.

To specifically identify the lipids secreted by the M14^KO^-brown adipocytes we applied comprehensive untargeted liquid chromatography-mass spectrometry (LC-MS) signaling lipidomics to five sets of samples. First, we collected plasma from overnight-fasted mice for evaluating circulating lipids (Figure 3A, left panel). In addition, we performed an *ex vivo* culture of the iBAT explant and collected conditioned Krebs solution as well as the iBAT tissue after a 2-hr culture at 37°C for examining BAT-secreted lipids (Lynes et al. 2017) (Figure 3A, left panel). Furthermore, considering iBAT is a heterogeneous tissue, we employed stable M14^KO^ hBAT cells to ensure the secreted factors are specifically from brown adipocytes. Thus, we differentiated hBAT cells, and collected conditioned culture media as well as cell pellets (Figure 3A, right panel).

**Figure 3.**
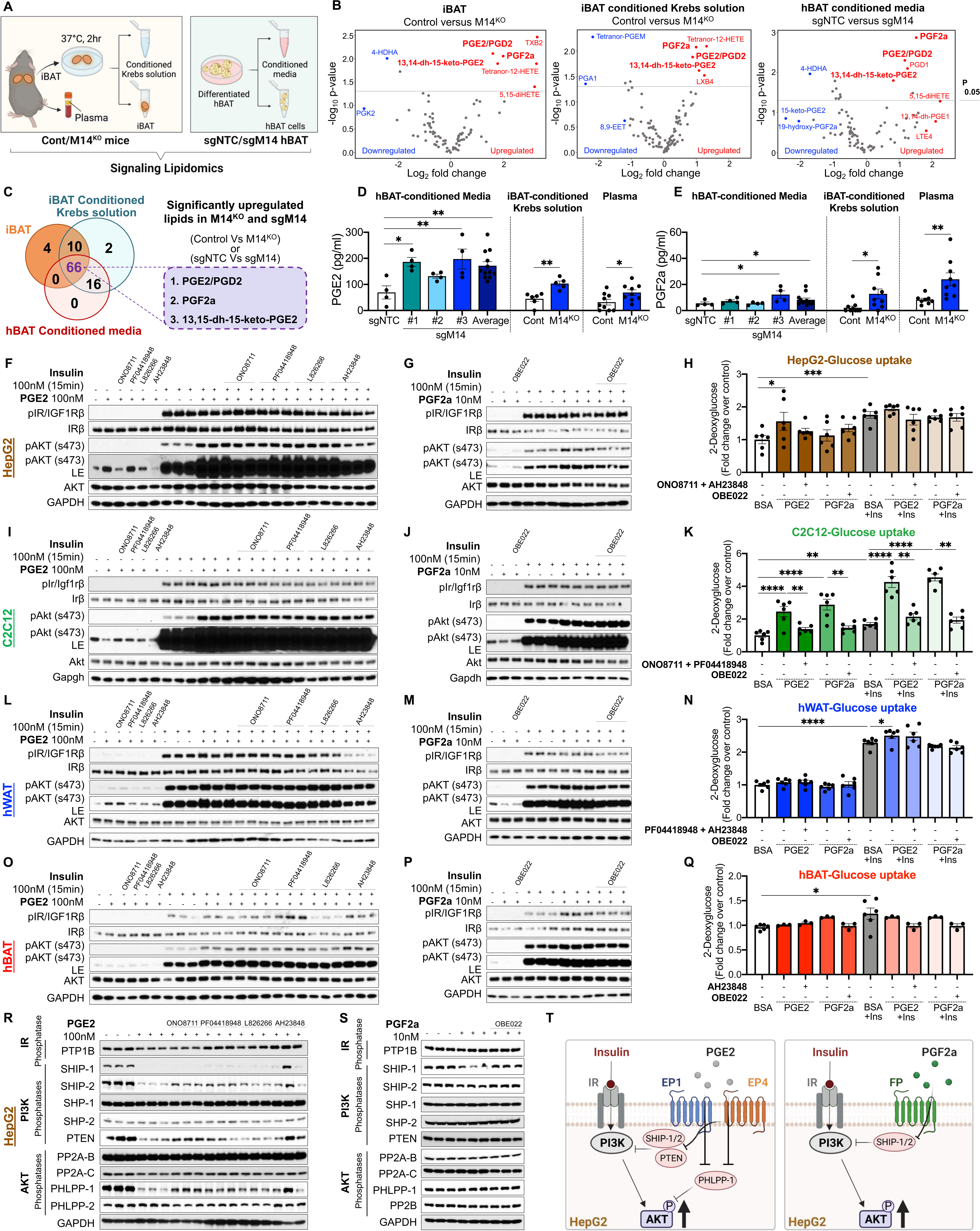
Ablation of Mettl14 Improves Insulin Sensitivity Primarily through Prostaglandin E2 and Prostaglandin F2a. (A) Schematic of sample preparation for untargeted signaling lipidomics. (B) Volcano plots of differentially down- (blue data points) and up-regulated (red data points) lipids in M14^KO^-iBAT, M14^KO^-iBAT conditioned Krebs solution, and sgM14-hBAT conditioned media identified by liquid chromatography mass spectrometric lipid analysis (n=9 for iBAT and iBAT conditioned Krebs solution, n=3-6 for hBAT conditioned media, log2 (fold change) threshold=2, *p* value threshold=0.05). (C) Venn diagram of all differentially abundant lipids in M14^KO^-iBAT, M14^KO^-iBAT conditioned Krebs solution, and sgM14-hBAT conditioned media. (n=9 for iBAT and iBAT conditioned Krebs solution, n=3-6 for hBAT conditioned media, log2 (fold change) threshold=2, *p* value threshold=0.05). (D and E) PGE2 (D) and PGF2a (E) concentrations in hBAT conditioned media, iBAT conditioned Krebs solution, and mouse plasma measured by ELISA assays. (F, I, L and O) Western blot of pIR/IGF1Rβ and pAKT_S473_ in HepG2 (F), C2C12 (I), hWAT (L), and hBAT (O) cells induced by PGE2 (100 nM) or insulin (100 nM) pre-treated with EP1/2/3/4 receptor antagonists ONO8711, PF04418948, L826266, and AH23848 (LE, longer exposure; n=3 biological replicates). (G, J, M, and P) Western blot of represented pIR/IGF1Rβ and pAKT_S473_ in HepG2 (G), C2C12 (J), hWAT (M), and hBAT (P) cells induced by PGF2a (10 nM) or insulin (100 nM) pre-treated with FP receptor Antagonist OBE022 (LE, longer exposure; n=3 biological replicates). (H, K, N, and Q) 2-Deoxyglucose uptake in in HepG2 (H), C2C12 (K), hWAT (N), and hBAT (Q) cells pretreated with PGE2 (100 nM for HepG2, C2C12, hWAT, and hBAT cells) or PGF2a (10 nM) and indicated antagonist(s) for overnight, followed by treating with insulin (100 nM) for 15 min (n=3 biological replicates). (R and S) Western blot of selected phosphatases in HepG2 cells pretreated with PGE2 (R) or PGF2a (S) in the presence of indicated antagonists (n=3 biological replicates). (T) Proposed model for mechanisms of action for PGE2 and PGF2a in HepG2 cells. Data are presented as mean ± SEM of biologically independent samples. *p < 0.05, **p < 0.01, ***p < 0.001 by two-tailed unpaired t-test (D, E, H, K, N, and Q). Also see Figure S3-S5, and Supplementary table 2.

Multivariate statistical models of the lipid profiling data were used to identify common lipids enriched in the five M14^KO^ sample types (Figures S3B-S3F). Comparative analyses of the individual signaling lipids between control and M14^KO^ samples (Figures 3B, S3G and S3H) led to the identification of 66 species that showed overlap among iBAT, iBAT conditioned Krebs solution, and hBAT conditioned media (Figure 3C). Besides, when intersecting iBAT, conditioned Krebs solution, and plasma samples, 75 lipids were commonly expressed among the three sets (Figures S3I). We noted that 73 lipids overlapped between conditioned hBAT media and hBAT cells (Figures S3J), while 82 lipids overlapped among conditioned hBAT media, conditioned Krebs solution, and plasma (Figures S3K). Importantly, 4 lipid species that belonged to the prostanoid family, namely prostaglandin E2 (PGE2), prostaglandin D2 (PGD2), prostaglandin F2 alpha (PGF2a), and 13,14-dihydro-15-keto-prostaglandin E2 (13,14-dh-15-keto-PGE2) were consistently significantly (*p < 0.05*) increased in a majority of the samples (M14^KO^ iBAT, M14^KO^ iBAT conditioned Krebs solution, and sgM14 hBAT conditioned media) (Figure 3C). Next, we validated this observation by confirming the upregulation of PGE2 and PGF2a in independent sets of mouse plasma, iBAT conditioned Krebs solution, and hBAT culture media using enzyme-linked immunosorbent assays (ELISA). Consistently, the absolute concentration of these 2 lipids were significantly higher in the M14^KO^ samples compared to controls (Figures 3D and 3E). These data suggest PGE2, PGD2, PGF2a, and 13,14-dh-15-keto-PGE2 are potential M14^KO^-BAT secreted lipokines.

To directly validate whether the candidate prostaglandins are responsible for the observed improvement in insulin sensitivity in peripheral metabolic cells, we performed a series of independent dose-response studies in a human liver cell line (HepG2) (Figure S4A), a differentiated mouse skeletal muscle cell line (C2C12) (Figure S4B), differentiated hWAT cell (Figure S4C), or differentiated hBAT cells (Figure S4D). Briefly, cells were treated with physiological (10, 100 nM), or supraphysiological concentrations (1000 nM) of PGE2, PGD2, PGF2a or 13, 14,dh-15-keto-PGE2 overnight. Cells were then washed and stimulated with 100 nM insulin for 15 mins (Figures S4A-S4D). Among the 4 candidate signaling lipids, PGE2 (100 nM) and PGF2a (10 nM), respectively, increased insulin stimulated pAKT_S473_ under both physiological and palmitate-induced insulin resistant conditions in HepG2 (Figure S4B), C2C12 (Figure S4D), hWAT (Figure S4F), or hBAT cells (Figure S4H) (results are summarized in supplementary table 2). These results point to the potency of physiological concentrations of PGE2 and PGF2a in improving insulin signaling in metabolic cells *in vitro*.

Next, we sought to identify the receptors and intracellular signaling networks that mediate the insulin sensitizing effects of PGE2 and PGF2a. Prostaglandins are known to exert their actions by acting on G-protein-coupled receptors (GPCRs) (Ricciotti and FitzGerald 2011). There are four GPCR designated subtypes (EP1, EP2, EP3, and EP4) that mediate the actions of PGE2 (Burkett, Doran, and Gannon 2023). Furthermore, the biological actions of PGF2a are mediated through binding with the prostaglandin F receptor (FP) (Woodward, Jones, and Narumiya 2011). We therefore hypothesized that PGE2 or PGF2a exert the insulin sensitizing effects via binding to these receptor(s). To directly test this hypothesis, we performed independent blocking experiments in HepG2, C2C12, differentiated hWAT or hBAT cells by pre-treating these cells with an antagonist of EP1(ONO8711) (Watanabe et al. 1999), EP2 (PF04418948) (af Forselles et al. 2011), EP3 (L826266) (Juteau et al. 2001), EP4 (AH23848) (Brittain et al. 1985), or FP (OBE022) (Pohl et al. 2018), receptor respectively. This was followed by treating the cells with PGE2 (100 nM) or PGF2a (10 nM) at concentrations determined from earlier experiments (Figures S4E-S4H). As a functional read out, we examined glucose uptake in HepG2, C2C12, hWAT, or hBAT cells. Similar to the treatment for assessing insulin signaling as described above, cells were pre-treated with specific receptor antagonist(s), followed by overnight treatment with PGE2 (100 nM), or PGF2a (10 nM) and an acute insulin stimulation (100 nM for 15 min) prior to assaying for 2-deoxy-glucose (2DG) uptake.

In HepG2 cells, 100 nM of PGE2 treatment induced pAKT_S473_ activation independent of insulin (Figures 3F, pAKT_S473_ longer exposure and S5A). Pre-treatment with EP1, EP3, or EP4 antagonists, namely ONO8711, L826266, or AH23848, blunted the PGE2-induced pAKT_S473_ even in the presence of insulin (Figures 3F and S5A). PGE2 synergistically increased pAKT_S473_ without influencing pIRβ/IGF1Rβ; furthermore, inhibition of EP1 or EP4 receptor by ONO8711 or AH23848 significantly abolished the synergistic effects of PGE2 on insulin (Figures 3F and S5A). On the other hand, PGF2a did not activate pAKT_S473_ *per se,* however, it increased insulin-stimulated pAKT_S473_ likely through actions on FP receptors in liver cells since OBE022 abolished the PGF2a-induced increase in pAKT_S473_ (Figures 3G and S5B). As expected, insulin induced an increase in glucose uptake in HepG2 cells (Figure 3H). Interestingly, PGE2 induced a 1.6-fold increase in glucose uptake at 100 nM, although this uptake was not as robust as the effect of insulin in HepG2 cells (Figure 3H).

In C2C12 cells, neither PGE2 nor PGF2a increased the basal pAkt_S473_ levels, however both prostaglandins elevated insulin-stimulated pAkt_S473_ (Figures 3H, 3I, S5C, and S5D). Strikingly, individual treatments with PGE2 (100 nM), or PGF2a (10 nM), led to a 2.6-fold or 3.2-fold induction of glucose uptake in C2C12 cells independent of insulin respectively, whereas insulin on its own induced a 2-fold increase in glucose uptake as expected (Figure 3K). Simultaneous blockage of EP1 and EP2 receptors abolished PGE2-induced glucose uptake. Similarly, the use of a PF receptor antagonist significantly decreased PGF2a-induced increase in glucose uptake in C2C12 cells (Figure 3K).

In differentiated hWAT cells, PGE2 but not PGF2a increased basal pAKT_S473_ levels, and both PGs increased insulin-stimulated pAKT_S473_ (Figures 3L, 3M, S5E, and S5F). Inhibition of EP2 or EP4 receptor, and FP receptor decreased PGE2-or PGF2a-induced effects, respectively. Conversely, blockage of EP1 or EP2 receptor partially decreased PGE2-induced effect while the FP antagonist, OBE022, did not significantly change PGF2a-induced effects (Figures 3L, 3M, S5E, and S5F). Besides, neither PGE2 nor PGF2a had an effect on glucose uptake in the absence of insulin, while PGE2 enhanced insulin-stimulated glucose uptake (Figure 3N).

In differentiated hBAT cells, both PGE2 and PGF2a increased insulin-stimulated pAKT_S473_ without directly impacting pAKT_S473_ (Figures 3O, 3P, S5G, and S5H). Interestingly, blockage of EP4 receptor showed a significant increase in PGE2-induced pAKT_S473_ (Figures 3O, and S5G). On the other hand, FP receptor antagonist completely abolished PGF2a-induced elevation in insulin-stimulated pAKT_S473_ (Figures 3P and S5H). Glucose uptake in hBAT cells was significantly stimulated by insulin, but was not altered in response to both PGs (Figure 3Q), suggesting that brown adipocytes are unlikely to be major target of PGs. These data together suggest that PGE2 and PGF2a act as secretory factors that exogenously control glucose uptake primarily in muscle cells, hepatocytes and white adipocytes.

Lastly, we investigated the molecular mechanism(s) activated by PGE2 or PGF2a to increase pAKT_S473_. Several mechanisms exist to attenuate, fine-tune, or terminate insulin signaling, both at the level of the receptor and downstream points in the cascade (Seely et al. 1996; Taniguchi, Emanuelli, and Kahn 2006). For example, phosphatases such as PH domain and Leucine rich repeat Protein Phosphatases (PHLPPs), SH2 domain-containing inositol phosphatase 1/2 (SHIP-1/SHIP-2), and Phosphatase and tensin homolog (PTEN) are known to negatively regulate AKT in the insulin signaling cascade (Virkamäki, Ueki, and Kahn 1999). We therefore hypothesized that PGE2 and PGF2a suppress the expression of specific negative regulators in the PI3K pathway and activities of pAKT. Using HepG2 cells as a model we observed that PGE2 decreased the protein abundance of PHLPP-1 which acts to dephosphorylate AKT; and SHIP-1/2 as well as PTEN, all of which negatively regulate the PI3K pathway. Blocking EP1 or EP4 receptors partially rescued the expression of these phosphatases (Figures 3R and S5I). PGF2a decreased the protein levels of SHIP-1 and SHIP-2, and this effect was reversed by blocking the FP receptor (Figures 3S and S5J).

Taken together, four signaling lipids, namely PGE2, PGD2, PGF2a, and 13,14-dh-15-keto-PGE2 are released by mouse brown adipocytes or human brown adipocytes that lack METTL14. Among these, PGE2 and PGF2a potentially act as primary endocrine factors mediating improved peripheral insulin sensitivity. For example, PGE2 stimulates pAKT_S473_ in the absence of insulin, and sensitizes insulin-stimulated pAKT_S473_ through distinct cell-specific EP receptors. In HepG2 cells, these actions are exerted by decreasing the expression of PHLPP-1, SHIP-1/2 or PTEN respectively. PGF2a acts by binding with FP receptor to negatively regulate SHIP-1 and SHIP-2 to increase insulin-stimulated pAKT_S473_ (Figure 3T).

### Exogenous administration of PGE2 and PGF2a Improves Insulin Sensitivity and Glucose Tolerance in DIO Mice

After observing improved insulin signaling and glucose uptake consequences of PGE2 and PGF2a treatment in *in vitro* metabolic cell models, we undertook long-term PGE2 and PGF2a administration studies to phenocopy the M14^KO^ mouse model. To this end, 13-week old LFD- or HFD-fed male mice were injected intra-peritoneally with 25 mg/kg PGE2+PGF2a (12.5 mg/kg PGE2 + 12.5 mg/kg PGF2a) or 50 mg/kg PGE2+PGF2a (25 mg/kg PGE2 + 25 mg/kg PGF2a) every other day for three weeks (Figure 4A). We monitored the activity, basal core body temperature and food intake as indicators of potential side effects induced by chronic PG administration. PG administration, at neither dose, impacted the activity (Figures S6A and S6B) or basal core body temperature (Figure S6C) in the fed LFD or HFD, pointing to the safety of PG administration in mice. Besides, combined PGE2+PGF2a injection at both doses did not significantly affect the food intake (Figure S6D) or body weight trajectories except for mice receiving 25 mg/kg PGE2+PGF2a challenged a HFD (Figure 4B). Furthermore, DIO mice injected with the low dose of PGE2+PGF2a had smaller body size, and decreased size and weights of iBAT, liver, iWAT and eWAT (Figures 4C and 4D); whereas these differences were not observed among the LFD-fed groups (Figures 4B, 4D and S6B). Histological analyses of the three adipose depots and liver tissues revealed healthier morphology in the PGE2+PGF2a-injected mice on HFD (Figures 4E-4J). Specifically, the PGs decreased HFD-induced hepatic steatosis (Figure 4E), immune cell infiltration and adipocyte hypertrophy in eWAT (Figures 4F) and iWAT (Figures 4G), and limited the adipocyte hypertrophy in iBAT (Figures 4H). However, no alterations were observed in the muscle (Figures 4I) or pancreas (Figures 4J). Interestingly, we also observed an induction of beige cells, containing multilocular lipid droplets, in the iWAT of mice injected with 50 mg/kg PGs (Figure 4G).

**Figure 4.**
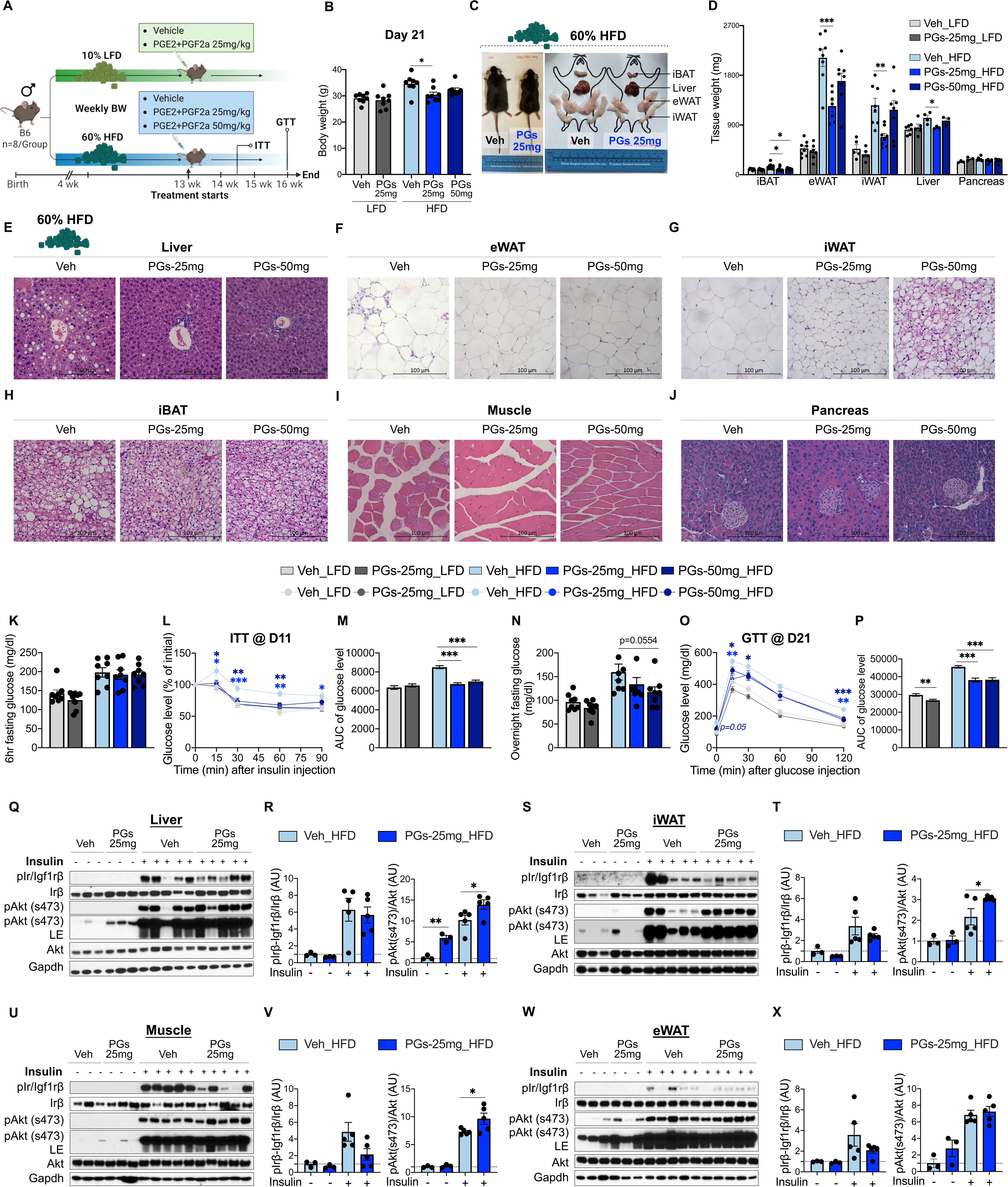
Administration of PGE2 and PGF2a Improves Insulin Sensitivity in HFD-fed Mice. (A) Overview of administration of PGE2+PGF2a on every other day by intraperitoneal injections of Vehicle (Saline), 25 mg/kg, or 50 mg/kg PGE2+PGF2a into 12-week old LFD-fed or HFD-fed mice for 21 days (n=8/group). (B) Body weight of vehicle-, 25 mg/kg PGE2+PGF2a-, or 50 mg/kg PGE2+PGF2a-injected LFD- or HFD-fed mice after 3-week injection (n=8/group). (C) Representative gross appearance of body size, adipose tissues, and liver in vehicle- or 25 mg/kg PGE2+PGF2a-treated mice. (D) Tissue weight of iBAT, iWAT, eWAT, liver, pancreas of vehicle-, 25 mg/kg PGE2+PGF2a-, or 50 mg/kg PGE2+PGF2a-injected LFD- or HFD-fed mice after 3-week injection. (E-J) Representative H&E staining in liver (E), eWAT (F), iWAT (G), iBAT (H), muscle (I), and pancreas (J) from vehicle-, 25 mg/kg PGE2+PGF2a-, or 50 mg/kg PGE2+PGF2a-injected HFD-fed mice. (K-P) 6hr-fasting glucose levels (K), intraperitoneal insulin tolerance tests (L and M), overnight-fasting glucose levels (N), and intraperitoneal glucose tolerance tests (O and P) of vehicle-, 25 mg/kg PGE2+PGF2a-, or 50 mg/kg PGE2+PGF2a-injected LFD- or HFD-fed mice. (Q-X) Representative western blot of pIr/Igf1rβ and pAkt_S473_ in liver (Q and R), iWAT (S and T), muscle (U and V), and eWAT (W and X) after 21-day injection of saline, or 25 mg/kg PGE2+PGF2a, followed by 1U of insulin injection via *vena cava*. All samples in each panel are biologically independent. Data are expressed as means ± SEM. *p < 0.05, **p < 0.01, ***p < 0.001 by Two-tailed unpaired t-test (B, D, K, M, N, P, R, T, V, X) and Two-way ANOVA (L and O). Also see Figures S6.

To uncouple the insulin sensitivity from the effects of body weight changes, we performed an insulin tolerance test (ITT) on day 11 post PGE2+PGF2a injection when body weights were similar between groups (Figure S6F). Treatment with PGE2+PGF2a did not influence LFD-fed mice which exhibited normal insulin sensitivity, while protecting DIO mice from HFD-induced insulin resistance at both doses (Figures 4K-4M). At the end of the experiment, PGE2+PGF2a (50 mg/kg) injection significantly lowered the fasting glucose in HFD-fed mice (Figure 4N), and both doses of PG decreased HFD-induced glucose intolerance (Figures 4O and 4P). Furthermore, treatment of DIO mice with PG (25 mg/kg) followed by assessment of insulin sensitivity after *vena cava* insulin injection showed increased pAkt_S473_ in the liver (Figures 4Q and 4R), iWAT (Figures 4S and 4T), and muscle (Figures 4U and 4V), without significantly altering insulin sensitivity in eWAT (Figures 4W and 4X).

In addition, to understand whether the global improvement in insulin sensitivity induced by PGs were dependent of the canonical UCP1-mediated thermogenesis of BAT, we performed indirect calorimetry after acute cold exposure. Regardless of improved cold tolerance induced by both doses of PGs in the DIO mice (Figure S6G), no differences were observed in energy expenditure (Figure S6H), O_2_ consumption (Figure S6I), or CO_2_ production (Figure S6J) between PGs and Veh groups fed with HFD after 6hr acute cold exposure. These results support the notion that M14^KO^-BAT secreted prostaglandins improve whole body metabolism independent of BAT thermogenesis.

Collectively, long-term treatment of DIO mice with PGE2 and PGF2a phenocopied the M14^KO^ mouse model and support the concept that M14^KO^-BAT improves metabolic health by endocrine effects of the PGs.

### Plasma Levels of PGE2 and PGF2a Are Negatively Associated with Obesity and Insulin Resistance in Humans

Since M14^KO^ mice and PGE2+PGF2a-treated mice were resistant to HFD-induced insulin resistance, we sought to examine the relevance of these findings in humans. We therefore tested the relationship between circulating levels of PGE2 or PGF2a and metabolic parameters in three independent cohorts of human subjects with a broad distribution of BMI and insulin sensitivity.

In human cohort 1, including both lean and obese individuals (sample information in Supplemental table 3) (Leiria et al. 2019), we observed that PGE2 levels were significantly decreased in overweight (25 kg/m^2^ ≤ BMI < 30 kg/m^2^) and obese subjects (BMI ≥ 30 kg/m^2^) compared to leans (BMI < 25 kg/m^2^) (Figures 5A). Next, Spearman’s correlation analyses identified a significantly negative correlation between plasma levels of PGE2 and BMI (Figure 5B), and HOMA-IR (Figure 5C). Besides, PGE2 presented a negative correlation with fasting plasma insulin (FPI), and leptin levels (Figures S7A). Similarly, PGF2a levels were significantly lower in overweight and obese subjects than leans (Figure 4D), and inversely correlated with BMI (Figure 4E), HOMA-IR (Figure 4F), FPI and leptin (Figure S7B). Altogether these data suggest that PGE2 and PGF2a levels in humans are inversely correlated with HOMA-IR, indicating that low levels of plasma PGs are associated with increased insulin resistance.

**Figure 5.**
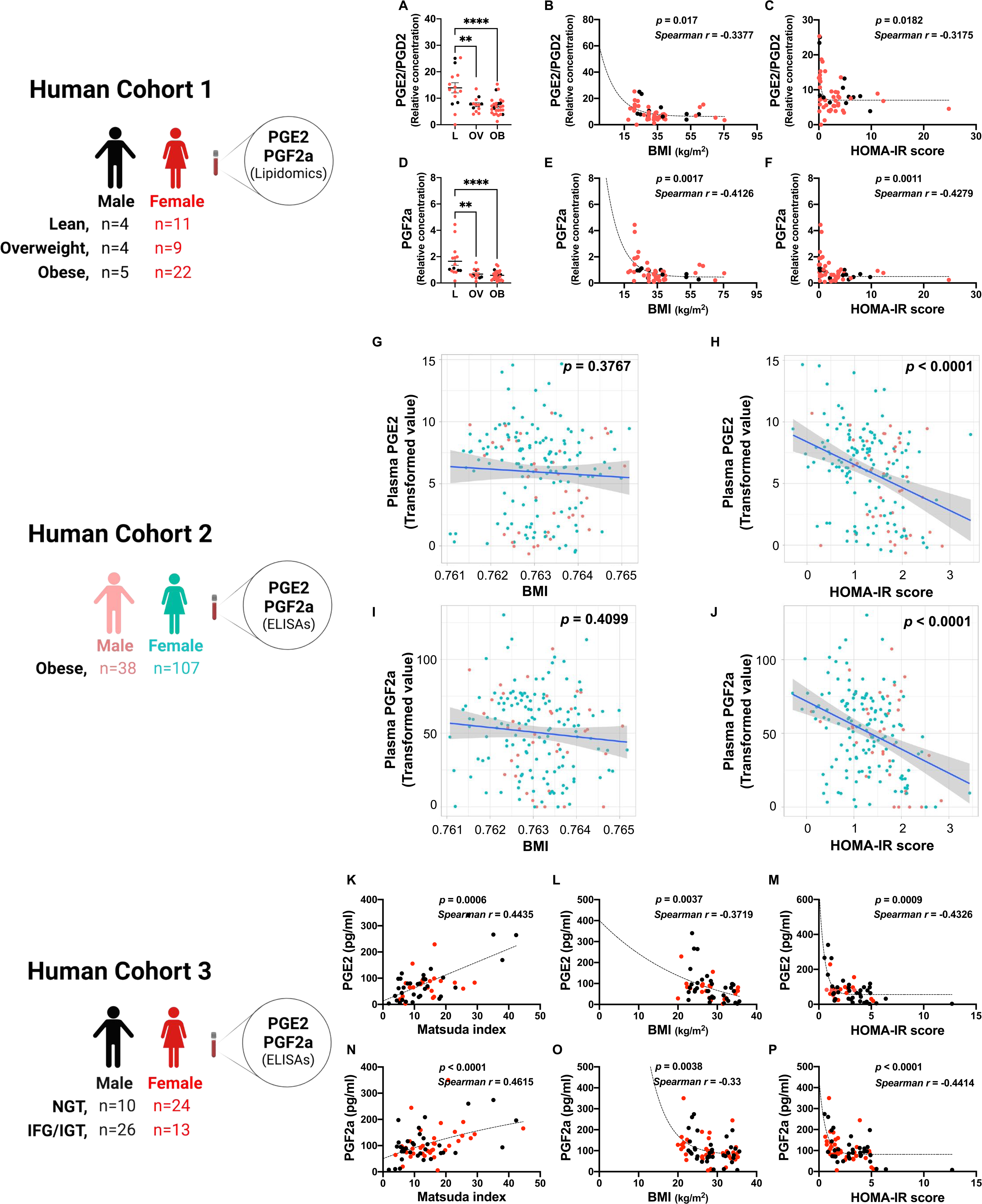
Plasma Levels of PGE2 and PGF2a Are Negatively Associated with Obesity and Insulin Resistance in Humans. (A) Plasma PGE2/PGD2 levels in lean (BMI < 25 kg/m^2^), overweight (BMI > 25 kg/m^2^, < 30 kg/m^2^), and obese (BMI > 30 kg/m^2^) subjects (n = 55). **p < 0.01, ***p < 0.001. Data are represented as mean ± SEM. L, lean; OV, overweight; OB, obese. (B) Spearman correlation between the plasma levels of PGE2/PGD2 and BMI. (C) Spearman correlation between the plasma levels of PGE2/PGD2 and insulin resistance measured by HOMA-IR. (D) Plasma PGF2a levels in lean (BMI < 25 kg/m^2^), overweight (BMI > 25 kg/m^2^, < 30 kg/m^2^), and obese (BMI > 30 kg/m^2^) subjects (n = 55). **p < 0.01, ***p < 0.001. Data are represented as mean ± SEM. (E) Spearman correlation between the plasma levels of PGF2a and BMI. (F) Spearman correlation between the plasma levels of PGF2a and insulin resistance measured by HOMA-IR. (G) Correlation between serum PGE2 levels and BMI in obese individuals (n=145). (H) Correlation between serum PGF2a levels and insulin resistance measured by HOMA-IR in obese individuals (n=145). (I) Correlation between serum PGE2 levels and BMI in obese individuals (n=94). (J) Correlation between serum PGF2a levels and insulin resistance measured by HOMA-IR in obese individuals (n=94). (K) Spearman correlation between the plasma levels of PGE2 and Matsuda index (n=55). (L) Spearman correlation between the plasma levels of PGE2 and BMI (n=55). (M) Spearman correlation between the plasma levels of PGE2 and insulin resistance measured by HOMA-IR (n=55). (N) Spearman correlation between the plasma levels of PGF2a and Matsuda index (n=73). (O) Spearman correlation between the plasma levels of PGF2a and BMI (n=73). (P) Spearman correlation between the plasma levels of PGF2a and insulin resistance measured by HOMA-IR (n=73). For human cohort 1, relative concentrations are used in (A-F) because the lipid quantification data were detected using non-targeted lipidomics. For human cohort 2, PGE2 and PGF2a concentrations were measured by ELISAs. Transformed values of metabolite and clinical variables were used. For human cohort 3, PGE2 and PGF2a levels were measured by ELISAs. Absolute concentrations were used. All samples in each panel are biologically independent. *p < 0.05, **p < 0.01, ***p < 0.001. Also see Figure S7, and Supplementary tables 3-5.

To further dissect the insulin sensitizing effects of the PGs independent of BMI, we next examined the association of circulating PGE2 or PGF2a levels with the metabolic parameters in human cohort 2, (also referred as Kuopio Obesity Surgery (KOBS) study) including only obese subjects (BMI ≥ 30) (De Jesus et al. 2020). The absolute levels of PGE2 and PGF2a were measured by ELISAs. Plasma PGE2 levels did not show significant correlations with BMI (Figure 5G), which allowed us to dissect the potential effects of PGs from body weight regulation in this human cohort. Consistent with human cohort 1, we also observed a negative correlation between PGE2 and HOMA-IR in human cohort 2 (Figures 5H). In addition, PGE2 showed negative correlations with fasting glucose, insulin, and triglycerides (Figures S7C). PGF2a showed a highly similar pattern of correlations with the parameters mentioned above (Figures 5I, 5J, and S7D).

Finally, we proceeded to determine the relationship between circulating levels of PGE2 and PGF2a levels with peripheral insulin sensitivity. In human cohort 3 (also referred as Stop Diabetes (StopDia) study) where Matsuda index, which represents both hepatic and peripheral tissue sensitivity to insulin is available, we found a significantly positive correlation between PGE2 and the Matsuda index (Figure 5K). Besides, plasma levels of PGE2 were found to negatively correlate with BMI (Figure 5L), HOMA-IR (Figure 5M), and FPI (Figure S7E), respectively. Similarly, PGF2a level was negatively correlated with these parameters (Figures 5N-P, and S7F). Overall, these findings suggest that PGE2 and PGF2a are linked to systemic metabolism and insulin sensitivity in humans.

### METTL14-deficiency Upregulates Pathways Related to Prostaglandin Synthesis in BAT

To answer the lingering question regarding the mechanism(s) by which METTL14 regulates prostaglandin synthesis in brown adipocytes we began by determining the transcriptional basis of prostaglandin synthesis following METTL14 ablation. We performed RNA sequencing (RNA-seq) and m^6^A-RNA immunoprecipitation sequencing (m^6^A-RIP-seq) in iBAT from control or M14^KO^ mice (Figure 6A). Principal component analysis (PCA) of both RNA-seq and m^6^A-seq data showed clear separation between control- and M14^KO^-iBAT (Figures 6B and 6C, respectively). RNA-seq identified 697 differentially expressed genes (DEGs) (p < 0.01) in M14^KO^-iBAT consisting of 324 upregulated and 373 downregulated genes (Figure 6D). Pathway and process enrichment analyses of the significantly upregulated genes revealed metabolism of lipids and phospholipid metabolic process among the top upregulated pathways as a consequence of M14 deficiency (Figures 6E and S8A). Consistently, pathways including response to insulin, adipogenesis, and glucose homeostasis were upregulated in M14^KO^-iBAT compared to controls. On the other hand, top-ranking ontologies for the downregulated genes were related to angiogenesis, cell cycle and mitosis (Figure S8B).

**Figure 6.**
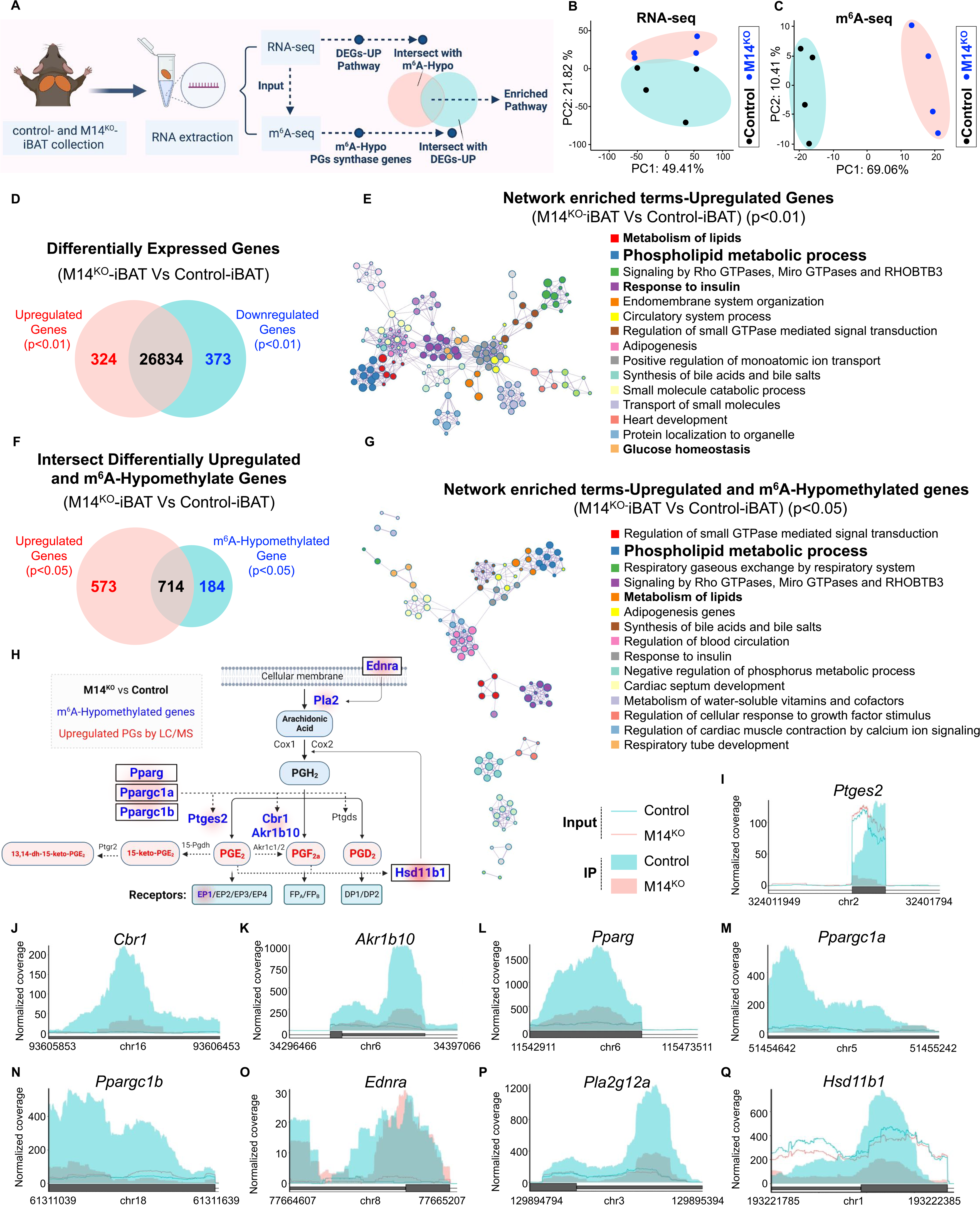
METTL14 Selectively Methylates Transcripts Encoding Prostaglandin Synthases and Their Regulators. (A) Schematic illustration of the RNA-seq and m^6^A-seq bioinformatic analyses strategy of iBAT samples from control and M14^KO^ mice. (B) PCA plot of RNA sequencing in controls (black dots, n=4 independent biological samples) and M14^KO^-iBAT (blue dots, n=4 independent biological samples). (C) PCA plot of m^6^A-seq in controls (black dots, n=4 independent biological samples) and M14^KO^-iBAT (blue dots, n=4 independent biological samples). (D) Venn diagram representation of the upregulated (red), downregulated (blue), and unchanged genes (black) of M14^KO^-iBAT compared to control-iBAT. Statistical analyses were performed using the Benjamin-Hochberg procedure and genes were filtered for p < 0.01. (E) Top 15 enriched GOs and pathways of upregulated genes in M14^KO^-iBAT versus control-iBAT. (F) Venn diagram representation of the intersection between upregulated genes (red) with m^6^A-hypomethylated genes (blue) in M14^KO^-iBAT versus control-iBAT. Genes were filtered for p < 0.05. (G) Functional enrichment of intersected genes in (F). (H) Representation of Prostaglandins biosynthesis pathway based on KEGG and Wikipathway annotations depicting several m^6^A hypomethylated genes (blue) and unchanged genes (black), and the upregulated prostaglandins suggested by LC/MS lipidomics (red) in M14^KO^-iBAT versus control-iBAT (genes filtered for p < 0.05). (I-Q) Coverage plots of m^6^A peaks in *Ptges2* (I), *Cbr1* (J), *Akr1b10* (K), *Pparg* (L), *Ppargc1a* (M), *Ppargc1b* (N),, *Ednra* (O), *Plasg12a* (P), and *Hsd11b1* (Q) genes in M14^KO^-iBAT versus control-iBAT. Plotted coverages are the median of the n replicates presented. All samples in each panel are biologically independent. Data are expressed as means ± SEM. *p < 0.05, **p < 0.01, ***p < 0.001. Also see Figure S8.

As previously reported (Dominissini et al. 2012; Meyer et al. 2012), the m^6^A peaks in both control and M14^KO^-iBAT were enriched at the start and stop codons and were characterized by the canonical GGACU motif (Figure S8C). To further test whether the transcriptomic alterations related to the phospholipid synthesis pathways were regulated by METTL14-mediated m^6^A decoration, we intersected the differentially upregulated (p < 0.05) and m^6^A-hypomethylated genes (p < 0.05) and obtained 714 transcripts (Figure 6F). Consistent with our RNA-seq data, this group of 714 genes was the most enriched for phospholipid metabolic process and metabolism of lipids pathways (Figures 6G and S8D).

Prostaglandin synthesis occurs when a cyclooxygenase enzyme (Cox1 and Cox2, also known as Ptgs1 and Ptgs2, respectively) converts arachidonic acid to an intermediate that is subsequently converted to an active prostanoid such as PGE2 and PGF2a. The biosynthesis of diverse PGs from arachidonic acid is directly enabled by various PG synthases such as the prostaglandin E Synthases (Ptges, Ptges2, Ptges3) but also indirectly regulated by several transcription regulators or transcriptional co-activators (Chambers et al. 2020; Bogacka et al. 2013). To further identify the target transcripts of METTL14 involved in the PG biosynthesis pathway, we analysed the m^6^A levels of the genes encoding prostaglandin synthases in control- and M14^KO^-iBAT. Several genes involved in the PG biosynthesis pathway were highly hypomethylated in the M14^KO^-iBAT (Figure 6H). More specifically, prostaglandin E synthase 2 (*Ptges2*) (Figure 6I), which encodes the synthase for PGE2 synthesis, Carbonyl reductase 1 (*Cbr1*) (Figure 6J) and aldo-keto reductase family 1 member B10 (*Akr1b10*) (Figure 6K), which are responsible for the production of PGF2a showed significantly lower m^6^A levels. Other hypomethylated genes in this pathway included peroxisome proliferator activated receptor gamma (*Pparg*) (Figure 6L), which promotes the synthesis/release of PGE2 (Bogacka et al. 2013), peroxisome proliferative activated receptor gamma coactivator 1 alpha (*Ppargc1a*) (Figure 6M), which promotes the expression of prostaglandin-endoperoxide synthase 1 (*Ptgs1*) (Chambers et al. 2020), and peroxisome proliferative activated receptor gamma coactivator 1 beta (*Ppargc1b*) (Figure 6N), which is a known regulator of several prostaglandin synthases. Among other hypomethylated genes in M14^KO^-iBAT were endothelin receptor type A (*Ednra*) (Figure 6O), phospholipase A2, group XIIA (*Pla2g12a*) (Figure 6P), which induce liberation of arachidonic acid from cellular membranes, as well as 11β-Hydroxysteroid dehydrogenase type 1(*Hsd11b1*) (Figure 6Q), which promotes conversion of arachidonic acid into prostaglandin H2 (PGH2) (Figure 6H). Thus, these three transcripts might contribute to the production of prostaglandins by providing additional substrates (i.e. arachidonic acid and PGH2). Consistent enriched pathways and genes were also observed in the sgM14-hBAT cells (Figure S8E and S8F). Overall, these results indicate deficiency of METTL14 directly and indirectly regulates the synthesis of prostaglandins in brown adipocytes.

### METTL14-mediated m^6^A Installation Promotes Decay of Prostaglandin Synthases and mRNAs of their Regulators in a YTHDF2/3-dependent Manner

We next sought to validate our sequencing outcome and investigate the molecular mechanism(s) underlying the METTL14-mediated m^6^A decoration of target transcripts in the PG synthesis pathway. First, qRT-PCR analyses confirmed the upregulation of *Pparg*, *Ppargc1a*, *Ppargc1b*, *Ptges2*, *Crb1,* and *Akr1b10* in the M14^KO^-iBAT (Figure 7A). Interestingly, although no alteration in the m^6^A levels was detected, we observed an upregulation of *15-Pgdh* and *Ptgds* gene expression in M14^KO^-iBAT (Figure S9A). The protein levels of Pparg, Pgc1a, Pgc1b, Ptges2, Crb1, and Akr1b10 were also significantly increased in the iBAT of M14^KO^ mice (Figures 7B and S9B). Consistently, gene expression (Figures 7D and S9C) and protein levels (Figures 7E and S9D) of PG synthase enzymes as well as their positive regulators were increased in METTL14 deficient hBAT cells.

**Figure 7.**
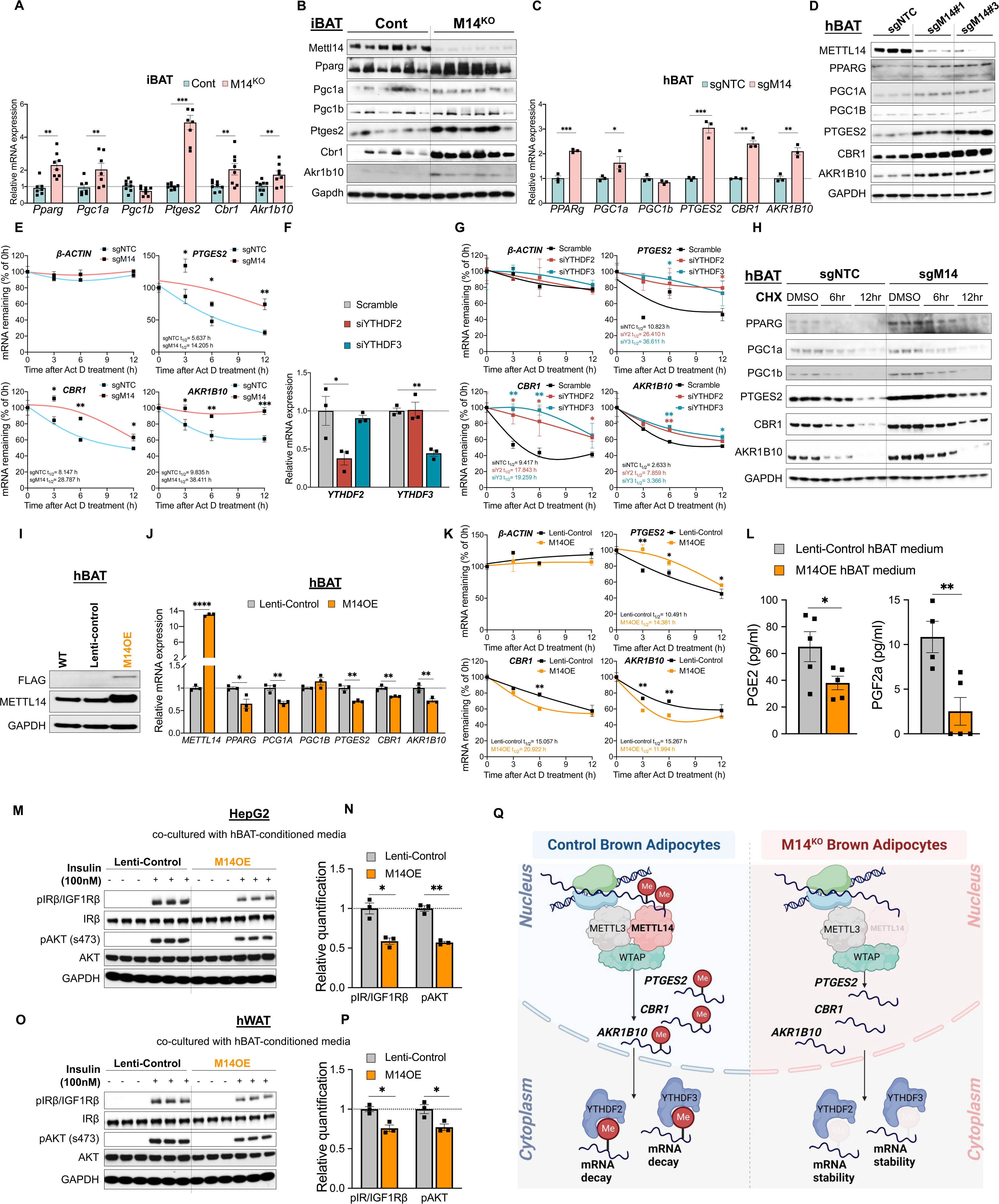
METTL14-mediated m^6^A Installation Promotes Decay of PGs synthases and Their Regulators mRNA in a YTHDF2/3-dependent Manner. (A) qRT-PCR analysis of the indicated mRNAs in the iBAT of male control and M14^KO^ mice. mRNA levels were normalized to β*-actin* (n = 8). (B) Western blot analysis of the indicated proteins in the iBAT of male control and M14^KO^ mice. Gapdh was used as a loading control (n = 6). (C) qRT-PCR analysis of the indicated mRNAs in the sgNTC and sgM14 hBAT cells. mRNA levels were normalized to β*-ACTIN* mRNA (n =3). (D) Western blot analysis of the indicated proteins in the the sgNTC and sgM14 hBAT cells. GAPDH was used as a loading control (n =3). (E) qRT-PCR analysis of the indicated mRNAs in differentiated sgNTC or sgM14 hBAT cells after a time-course treatment with 100 ug/mL Actinomycin D (Act D). mRNA levels were normalized to β*-ACTIN* mRNA (n = 3). (F) qRT-PCR analysis of *YTHDF2* and *YTHDF3* mRNAs in differentiated wildtype hBAT cells transfected with siNTC/siYTHDF2/siYTHDF3 siRNA. mRNA levels were normalized to β*-ACTIN* mRNA (n = 3). (G) qRT-PCR analysis of the β*-ACTIN*, *PTGES2*, *CBR1*, and *AKR1B10* mRNA in differentiated wildtype hBAT cells transfected with siNTC/siYTHDF2/siYTHDF3 siRNA and treated with 100 ug/mL Act D for the indicated time. *PTGES2*, *CBR1*, and *AKR1B10* mRNA levels were normalized to β*-ACTIN* (n = 3). (H) Protein stability of indicated proteins in differentiated sgNTC or sgM14 hBAT cells incubated with 100 ug/mL Cycloheximide (CHX) for the indicated time. GAPDH was used as a loading control (n = 3). (I) Western bolt analysis for validating of M14 overexpression in hBAT cells. (J) qRT-PCR analysis of indicated mRNAs in the control and M14OE hBAT cells (n = 3). (K) qRT-PCR analysis of the indicated mRNAs in differentiated control or M14OE hBAT cells after a time-course treatment with 100 ug/mL Actinomycin D (Act D). mRNA levels were normalized to β*-ACTIN* mRNA (n = 3). (L) ELISA analysis of PGE2 and PGF2a levels in the culture media of differentiated control and M14OE hBAT cells (n = 5). (M and N) Western blot analysis (M) and quantification (N) of pIR/IGF1Rβ and pAKT_S473_ in HepG2 cells co-cultured with conditioned medium from control or M14OE hBAT cells (n = 3). (O and P) Western blot analysis (O) and quantification (P) of pIR/IGF1Rβ and pAKT_S473_ in differentiated hWAT cells co-cultured with conditioned medium from control or M14OE hBAT cells (n = 3). (Q) A proposed model for the molecular mechanism of action that METTL14-mediated m^6^A installation destabilizes transcripts encoding prostaglandins synthases. All samples in each panel are biologically independent. Data are expressed as means ± SEM. *p < 0.05, **p < 0.01, ***p < 0.001 by Two-tailed unpaired t-test (A, C, J, L, N, P) and Two-way ANOVA (E, G, K). Also see Figure S9.

We then took advantage of the hBAT cell model to conduct mechanistic studies. First, we examined the effects of METTL14-deficiency on the stability of candidate target mRNAs following treatment with actinomycin D (Act D), an inhibitor of transcription, in hBAT cells. METTL14-deficiency did not significantly affect the stability of β*-ACTIN* mRNA, however, it potently suppressed degradation of *PTGES2*, *CBR1*, and *AKR1B10* mRNAs in the differentiated hBAT cells (Figure 7E). These data suggest that METTL14-mediated m^6^A methylation regulates target mRNA levels by impacting their stability.

To confirm this hypothesis, we performed siRNA-mediated knockdown of the m^6^A reader proteins YTHDF2 and YTHDF3 (Figure 7F), which are important for mRNA stability (Shi et al. 2017; X. Wang, Lu, et al. 2014), and examined the expression and stability of *PTGES2*, *CBR1*, and *AKR1B10* mRNAs after ActD treatment in wild type hBAT cells. Knockdown of either YTHDF2 or YTHDF3 followed by ActD treatment did not affect the stability of β*-ACTIN* while significantly increasing stability of *PTGES2*, *CBR1*, and *AKR1B10* mRNAs (Figure 7G).

Next, we investigated the impact of METTL14-deficiency on the translation of target mRNAs by performing a chase experiment where global protein synthesis is blocked using the translation elongation inhibitor cycloheximide (CHX). Ablation of METTL14 did not affect the protein expression of GAPDH during the 12-hr treatment likely due to its long half-life (Figures 7H and S9E). Furthermore, translational efficiency of *PPARG*, *PGC1a*, *PGC1B*, *PTGES2*, *CBR1*, and *AKR1B10* mRNAs was not significantly affected by METTL14-deficiency in the hBAT cells, as indicated by similar degradation of these mRNAs after a time-course treatment with CHX (Figures 7H and S9E). These results support our hypothesis that METTL14 primarily regulates the transcriptional process rather than translational efficiency.

Finally, to validate the involvement of METTL14 in regulating the mRNA stability of PG synthase genes, we asked whether the converse, i.e. upregulating METTL14 accelerates decay of the target genes. METTL14 overexpression (M14OE) was induced by lentiviral vector and validated by Western blot and qRT-PCR analyses (Figures 7I and 7J, respectively). M14OE did not significantly change the adipogenic capacity of hBAT cells as indicated by Oil Red O staining (Figure S9F), however, it decreased the gene expression of *PPARG*, *PGC1a*, *PGC1B*, *PTGES2*, *CBR1*, and *AKR1B10* (Figure 7J). In addition, M14OE promoted the degradation of *PTGES2*, *CBR1*, and *AKR1B10* mRNAs in hBAT cells after ActD treatment (Figure 7K). Consistently, the secretion of both PGE2 and PGF2a were significantly lower in the M14OE-hBAT compared to control hBAT cells as assessed by ELISA (Figure 7L). Furthermore, pre-treatment of HepG2 cells (Figures 7M and 7N) or hWAT cells (Figures 7O and 7P) with M14OE-hBAT conditioned media before insulin stimulation led to the downregulation of pIR/IGF1Rβ and pAKT_S473_ levels in HepG2 and differentiated hWAT cells compared to cells co-cultured with control lentivirus.

Taken together, we have uncovered a unique mechanism regulating systemic insulin sensitivity in peripheral metabolic tissues. Our data suggests that METTL14 selectively methylates transcripts encoding PGE2 and PGF2a synthases thereby promoting the decay of these target genes, which results in a negative regulation of PGE2, PGD2, PGF2a and 13,15-dh-15-keto-PGE2 production in brown adipocytes (Figure 7Q).

## Discussion

Insulin resistance, which is characterized by reduced sensitivity to insulin in the liver, muscle, and adipose tissues, is known to be associated with glucose intolerance and type 2 diabetes (T2D). The activity of brown adipose tissue has been found to negatively correlate with body mass index (BMI) (Cypess et al. 2009; van Marken Lichtenbelt et al. 2009) while being positively associated with insulin sensitivity (van Marken Lichtenbelt et al. 2009). Therefore, targeting BAT presents one potential approach for the prevention and treatment of obesity and insulin resistance (Czech 2020; Shamsi, Wang, and Tseng 2021).

BAT secretes various bioactive factors that can either directly modulate brown adipocyte insulin sensitivity and/or importantly, act as endocrine factors to regulate insulin sensitivity in peripheral metabolic tissues independent of canonical thermogenesis. Some of these factors include proteins (e.g., Nrg4 (G.X. Wang, Zhao, et al. 2014), FGF21 (Keipert et al. 2020), FGF6 and FGF9 (Shamsi et al. 2020), signaling lipids or so-called lipokines such as 12,13-diHOME (Lynes et al. 2017; Stanford et al. 2018), omega-3 oxylipin 12-HETE (Leiria et al. 2019). Exosomal microRNAs have also been reported to regulate systemic metabolism by inter-organ crosstalk (Thomou et al. 2017; Zhao et al. 2022; Crewe and Scherer 2022). However, most of these factors have been reported to be released in response to cold stimulation (Leiria et al. 2019; Lynes et al. 2017; Shamsi et al. 2020; Sugimoto et al. 2022; Sustarsic et al. 2018), browning inducers (Whitehead et al. 2021), or exercise (Stanford et al. 2018)). For the first time to our knowledge, we describe a unique mechanism by which METTL14-mediated m^6^A mRNA methylation negatively regulates BAT prostaglandin synthesis.

M14^KO^ mice are protected from diet-induced obesity and show consistent improvement in insulin sensitivity and glucose tolerance regardless of gender or diet. These phenotypes are independent of body weight reduction and thermogenic capacity of BAT. A significant increase in insulin signaling is observed in key peripheral metabolic tissues in M14^KO^ mice. Interestingly, despite increased insulin signaling in the M14^KO^-iBAT, the abundance of UCP1 is decreased. This finding along with unaltered energy expenditure suggests the improved insulin sensitivity is mediated via an UCP1-independent mechanism. Given the fact that the anti-obesity effects of UCP1-dependent thermogenesis is supported by a proof-of-principle study in UCP1-knockout mice maintained at thermoneutrality (Feldmann et al. 2009); and that obese humans express minor amounts of UCP1 and most do not live in thermoneutral conditions (Kajimura and Spiegelman 2020), UCP1-independent mechanism(s) may have significant translational relevance. Furthermore, METTL14 deficiency induces non-cell-autonomous effects in terms of insulin sensitivity, as indicated by the *in vitro* co-culture experiments. All these observations strongly support the inter-organ communication hypothesis.

By intersecting iBAT, iBAT-conditioned Krebs solution, and hBAT-conditioned media lipodomics, we identified PGE2, PGD2, PGF2a, and 13,14-dh-15-keto-PGE2 as the top upregulated lipokines in M14^KO^ models. Besides, plasma measurements by ELISA suggested significantly higher absolute levels of both PGF2a and PGE2 in the circulation. The availability of these lipids in the circulation suggests that they can regulate distant metabolic tissues and/or cells, resulting in systemic effects that can influence insulin sensitivity and glucose metabolism. A previous study has also reported that prostaglandin I2 (PGI2) contributes to energy expenditure, and protects mice from diet induced obesity through PGI2/Ptgir/PPARγ signaling (Vegiopoulos et al. 2010). Although both PGE2 and PGF2a have been recognized as factors that trigger pro-inflammatory responses (Ricciotti and FitzGerald 2011) and despite conflicting evidence on prostaglandins and insulin sensitivity (Inazumi et al. 2020; Shamsi et al. 2020; Yasui et al. 2015; Vegiopoulos et al. 2010), our data strongly suggest that PGE2 and PGF2a are *de facto* M14^KO^-BAT secreted insulin sensitizers.

Furthermore, it is likely that the effects of PGE2 or PGF2a vary among tissues/cells depending on distribution of their receptors. Thus, it is crucial to identify the specific receptors to establish the tissue-specific contributions downstream of both lipids. PGE2 and PGF2a act through distinct receptors to regulate different cell signaling pathways to induce distinct actions. For example, PGE2 binds with E-Prostanoid (EP1-4) receptors while PGF2a exerts its biological effects by binding solely with the prostaglandin F receptor. The availability of receptor-specific antagonists allowed us to identify the receptor(s) responsible for the insulin-sensitizing actions of PGE2 versus PGF2a in HepG2, hWAT, C2C12, and hBAT cells, respectively. Inhibition of the corresponding receptor(s) in C2C12 and hWAT cells attenuated the glucose uptake promoted by PGE2/PGF2a. Importantly, we provide a mechanism by which PGE2 or PGF2a increases pAKT_S473_ signaling. PGE2 effectively decreases the negative regulators of the PI3K pathway and phosphatases of AKT in HepG2 cells, while PGF2a only reduces the expression of phosphatases of PI3K. These observations are consistent with the insulin-independent and insulin-dependent activation of pAKT_S473_ induced/promoted by PGE2 or PGF2a. The inhibition of phosphatases is dependent on the receptors, supporting the notion that PGE2 and PGF2a exert their functions by binding to specific G protein-coupled receptor(s) (see summary in Figure 3T).

The beneficial phenotypes observed in M14^KO^ mice can be recapitulated through chronic treatment with PGE2 and PGF2a, which mimics the physiological conditions present in the circulation of the M14^KO^ mouse model. Adipocyte-derived PGE2 has been reported to indirectly enhance systemic insulin sensitivity by promoting regulatory T cell proliferation (C. Wang et al. 2022). PGE2 has been shown to enhance M2 macrophages, whose activation is thought to protect adipocytes from insulin resistance (Luan et al. 2015). In the present study, systemic treatment with PGE2 and PGF2a directly improves insulin sensitivity of multiple metabolic tissues, including WATs in the DIO mice. Activating beige cells leads to increased glucose uptake, fatty acid oxidation and lipolysis, resulting in improved insulin sensitivity (Cheng et al. 2021). Interestingly, a beige phenotype was observe in the iWATs of M14^KO^ mice and PGE2+PGF2a treated mice, suggesting the possibility that PGE2, PGD2, 13,14-dh-15-keto-PGE2, and/or PGF2a are browning inducers. Indeed, previous work has suggested that PGE2 interacts with PPARγ to divert pre-adipocyte differentiation into beige adipocytes (García-Alonso et al. 2013; García-Alonso and Clària 2014), and PGD2 has been demonstrated to promote beige cell proliferation (Abe et al. 2022). To the best of our knowledge, this is the first evidence showing the browning inducing effect of PGF2a and 13,14-dh-15-keto-PGE2. However, future studies should confirm this observation and address the mechanism(s) by which these two PGs promote the process of white fat-browning.

Our observations confirm the existence of an endocrine response in iWAT of M14^KO^ mice, mainly triggered by PGE2 and/or PGF2a. Additionally, despite decreased expression of UCP1 in their iBAT, M14^KO^ mice showed increased tolerance to acute cold exposure and unchanged energy expenditure. This suggests a compensatory effect due to induction of beiging in iWAT that contributes to the observed phenotype. Importantly, the translational relevance of our *in vitro* and animal studies is demonstrated by the inverse correlations observed between circulating PGE2 or PGF2a levels and HOMA-IR in three independent human populations. These findings provide strong evidence supporting the hypothesis that the improvement in insulin sensitivity resulting from M14 deficiency is linked to increased levels of PGE2 and PGF2a.

The transcriptome and m^6^A methylome analyses indicate that the ablation of METTL14 leads to an upregulation in the metabolism of lipids and the phospholipid metabolism pathway. However, a key question remains: how does METTL14-mediated m^6^A regulation specifically affect the biosynthesis of prostaglandins in brown adipocytes? The biosynthesis of prostaglandins involves the sequential oxygenation of arachidonic acid by cyclooxygenases (COX1 and COX2) to form PGH2. Specific PGs are then formed through the actions of different terminal PG synthases on PGH2 (Figure 6H).

It is worth noting that several transcription factors and coactivators, such as PPARG, PGC1a, and PGC1b, can each regulate the activity of terminal PG synthases. Therefore, we focused specifically on the genes encoding PG synthases and their regulators. Our analyses revealed that the *Ednra*, *Pla2g12a*, and *Hsd11b1* genes were hypomethylated and upregulated in the M14^KO^-iBAT. Ednra and Pla2g12a enzymes are responsible for promoting the synthesis of arachidonic acid, while the Hsd11b1 enzyme-promoted regeneration of COX2 contributes to the production of PGH2 (Gong, Morris, and Brem 2008). It is likely that more substrates (arachidonic acid and PGH2) are released/converted for the subsequent conversion into specific PGs. Absence of METTL14 results in the upregulation of enzymes involved in the conversion of PGE2, PGD2, PGF2a, and 13,14-dh-15-keto-PGE2 in brown adipocytes through direct and indirect regulation. First, the transcripts encoding enzymes Ptges2 (PGE2 synthase), Cbr1, and Akr1b10 (PGF2a synthases) were significantly upregulated and m^6^A hypomethylated, suggesting direct regulation by METTL14. Second, transcription factors/coactivators *Pparg* and *Pgc1a* were also upregulated and m^6^A hypomethylated, potentially leading to the indirect upregulation of genes encoding Ptgds (PGD2 synthase) and 15-Pgdh (13,14-dh-15-keto-PGE2 intermediate synthase) proteins (Bogacka et al. 2013; Chambers et al. 2020). Overall, these findings suggest that METTL14-deficiency directly and indirectly upregulates the expression of enzymes involved in prostaglandin conversion in brown adipocytes.

Our mechanistic studies revealed that METTL14-deficiency leads to decreased m^6^A decoration in target transcripts, which does not alter their translation. Instead, downregulation of m^6^A protects *PTGES2*, *CBR1*, and *AKR1B10* from mRNA decay therefore increasing their expression at transcriptional levels. This notion is supported by the observation that downregulation of m^6^A readers YTHDF2 and YTHDF3, which are known to mediate the mRNA decay, enhance the stability of *PTGES2*, *CBR1*, and *AKR1B10* mRNA (Figure 7Q). Inversely, overexpression of METTL14 promotes decay of *PPARG*, *PGC1A*, *PTGES2*, *CBR1*, and *AKR1B10*, decreases the expression of their gene expression and the production of PGE2 and PGF2a in differentiated hBAT cells.

In conclusion, this study reveals a unique role of m^6^A modification in the regulation of BAT secretory function. The BAT-secreted prostaglandin action represents an unexpected lipid-induced signaling pathway with beneficial metabolic effects in multiple key metabolic tissues. These data provide a potential therapeutic strategy for the prevention and treatment of insulin resistance.

## Limitations of study

There are some limitations to this work. 1) BAT-secreted factors include proteins, lipids and exosomal miRNAs. In this study, we ruled out a major regulatory role of proteins by denaturing the conditioned media, however, we cannot completely exclude potential regulation by miRNAs. Indeed, m^6^A has been reported to regulate miRNA biosynthesis by participating in the processing of pre-miRNAs or splicing of pre-miRNAs, and our RNA-seq and m^6^A-seq suggested many genes being regulated by miRNAs. Therefore, a future study should investigate the miRNA profile in the M14^KO^-BAT and its subsequent role in the regulation of insulin sensitivity. 2) While PGE2 and/or PGF2a treatment protected mice from HFD-induced gain in body weight and corrected insulin resistance and glucose intolerance, it is important to note that future studies should benchmark the efficacy of PGE2 and PGF2a to other anti-diabetic drugs that have been reported to have high efficacy in mice, including thiazolidinediones. 3) Besides, both our M14^KO^ mice and the PGs-treated mice were less prone to diet-induced obesity, while their energy expenditure and food intake are comparable with controls. Given the fact that the UCP1^KO^ mice are resistant to DIO via secreting energy-storing metabolites in urine (Keipert et al. 2020), and M14^KO^-BAT has downregulated UCP1 expression, additional studies are warranted to explore whether these mice secrete more energy into their feces and/or urine.

## Data and code availability

- All data reported in this paper will be shared by the lead contact upon request.
- This paper does not report original code.
- Any additional information required to reanalyze the data reported in this paper is available from the lead contact upon request.

## Acknowledgments

The authors thank Joslin Animal Physiology Core, Joslin Bioinformatics Core (P30 DK36836). We would like to thank Stephane Gesta PhD, Michael Kiebish PhD, and Valerie Bussberg (BERG) for technical assistance with untargeted lipidomics. This work is supported by NIH grants R01 DK67536 (R.N.K.), UC4 DK116278 (R.N.K. and C.H.), and RM1 HG008935 (C.H.). R.N.K. acknowledges support from the Margaret A. Congleton Endowed Chair and C.H. is a Howard Hughes Medical Institute Investigator. D.F.D.J. acknowledges support by Mary K. Iacocca Junior Postdoctoral Fellowship and American Diabetes Association grant #7-21-PDF-140. Human Cohort 2: Kuopio Obesity Surgery Study (PI: J.P.) was supported by the Finnish Diabetes Research Foundation, Kuopio University Hospital Project grant (EVO/VTR grants 2005–2021), the Academy of Finland grant (Contract no. 138006), and the Finnish Cultural Foundation. Human Cohort 3: STOP DIABETES - from knowledge to solutions project was funded by the Strategic Research Council at the Academy of Finland in 2016–2019 (303537) and by the Academy of Finland 2020–2023 (332465), by the Novo Nordisk Foundation 2018–2020 (33980 and 63753), and by the Finnish Diabetes Research foundation.

## Author Contributions

L.X. and D.F.D.J. conceived the study, designed and performed experiments, analyzed the data, assembled figures, and wrote the manuscript. C.W.J. performed RNA/m^6^A-seq. and m^6^A LC-MS/MS experiments. J.B.W. assisted with m^6^A-seq. and m^6^A LC-MS/MS experiments. J.H. performed immunohistochemistry. A.D.F. assisted with animal experiments. T.T. performed correlation analyses for human cohort 1. C.C. provided qRT-PCR data of human BAT tissues. V.M. contributed to collecting plasma and other clinical data of human cohort 2. S.M.M. contributed to collecting plasma and other clinical data of human cohort 3. S.Y.W. generated METTL14-OE and control lentiviral vectors. I.O. assisted with animal experiments. M.B. provided human plasma samples for human cohort 1 and edited the manuscript. A.M.C. contributed to conceptual discussions and provided human brown adipose tissues. J.P.I. was responsible for the human cohort 2 as PI, and contributed to conceptual discussions. Y.H.T. contributed to conceptual discussions, shared lipidomics data of human cohort 1, and provided immortalized human white/brown pre-adipocytes and protocols for culture and differentiation. C.H. contributed to conceptual discussions and designed the experiments for RNA-seq and m^6^A-seq. R.N.K. conceived the study, designed the experiments, supervised the project, and wrote the manuscript. All the authors have reviewed, commented on, and edited the manuscript.

## Tables with titles and legends

**Supplementary Table 1**. related to Figure 1. Clinical characteristics of the molecular, cellular, and genetic characterization of human adipose tissue and its role in metabolism human cohort.

**Supplementary Table 2**. related to Figure 3. pAKT_S473_ increasing potency of candidate prostaglandins in HepG2, C2C12, hWAT, and hBAT.

Notes: + indicates positive effects on pAKT_S473_, − indicates no or negative effects on pAKT_S473_.

**Supplementary Table 3**. related to Figure 5. Clinical characteristics of Human cohort 1 study.

**Supplementary Table 4**. related to Figure 5. Clinical characteristics of the Kuopio Obesity Surgery (KOBS, Human cohort 2) study.

**Supplementary Table 5**. related to Figure 5. Clinical characteristics of the Stop Diabetes(StopDia, Human cohort 3) study.

**Supplementary Table 6**. Primers used for qRT-PCR.

## Methods

### EXPERIMENTAL MODEL AND SUBJECT DETAILS

#### Mouse models and treatment

Mettl14^fl/fl^ mice on a C57BL/6N background (De Jesus et al. 2019) were crossed with UCP1^CRE^ strain (stock no.024670, the Jackson Laboratory) mice, which do not harbor the nicotinamide nucleotide transhydrogenase (Nnt) mutation, to generate the knockouts. For studies on diet-induced obesity and long-term insulin resistance, control and M14^KO^ mice were fed either 10% fat LFD or a conventional 60% fat HFD for 12 weeks, starting at 4 weeks of age.

To examine tissue-specific insulin sensitivity, control and M14^KO^ mice were fasted for 16 hours. After being anesthetized, 1 U of insulin was injected directly into the *vena cava* (Michael et al. 2000). Tissues were harvested at the indicated time points as shown in Figure 2A.

#### Acute cold exposure

For studies on cold tolerance tests, control and M14^KO^ mice were caged individually at an ambient temperature of 5°C for 1 to 6 hours in temperature-controlled diurnal incubators (Caron Products and Services) with *ad lib* access to water and food. Rectal temperature was measured at 1h intervals for core body temperature. For chronic cold exposure, 10-week mice were first exposed to intermittent cold exposure (12 h at 4°C in the light cycle and 12h at room temperature (22°C)) for 3 days followed by 4 days of continued cold exposure at 4°C. Intraperitoneal insulin tolerance test (IPITT) and Intraperitoneal glucose tolerance test (IPGTT) were performed on day 1 and day 4 post chronic cold exposure.

#### Chronic treatment with prostaglandins

For chronic treatment with PGs, mice were mock injected with saline for three days prior to administering PGE2 and PGF2a to prevent stress-induced weight loss. Mice were then injected via the intraperitoneal route with vehicle (saline), or 12.5 mg/kg of PGE2 and PGF2a, or 25 mg/kg of PGE2 and PGF2a every other day for 3 weeks. Body weight was measured weekly. IPITT and IPGTT were performed on day 11 and day 21 post-treatment. Indirect calorimetry was performed on day 16 post-treatment after acute cold exposure. At the end of the experiment, tissues were collected after *vena cava* insulin injection as described before in mouse models and treatment section.

#### Studies on molecular, cellular, and genetic characterization of human adipose tissue and its role in metabolism

Brown adipose tissue samples from 13 adult (6 males and 7 females) participants were selected from a cohort of healthy volunteers and surgical patients known to have brown adipose tissue, identified by PET/CT (ClinicalTrials.gov Identifier: NCT02692885). Brown adipose tissues were collected from different depots including retroperitoneal, deep abdominal, deep periadrenal, and deep retroperitoneal depots. Samples were divided into lean (BMI< 30) or obese (BMI ≥ 30) groups. Donor information is summarized in Supplementary Table 1. qRT-PCR was performed to detect the gene expression of m^6^A writers, erases, and readers.

#### Studies related to lean and obese human subjects (human cohort 1)

As described previously (Leiria et al. 2019), a group of 55 people were selected from the Leipzig obesity biobank to represent a wide range of BMI and parameters of glucose metabolism. This group consisted of 42 women and 13 men with BMI ranging from 17.5 to 75.4 kg/m^2^. The group was divided into 3 subgroups: lean (15 individuals), overweight (13 individuals), and obese (27 individuals). In the lean subgroup, all individuals were normal glucose tolerant, while in the overweight subgroup, 10 were normal glucose tolerant and 3 had T2D, and in the obese subgroup, 20 were normal glucose tolerant and 7 had T2D. Patient information is summarized in Supplementary table 3. The collection of human biomaterial, serum analyses, and phenotyping was approved by the ethics committee of the University of Leipzig (approval numbers: 159-12-21052012 and 017-12-23012012) and all individuals provided written informed consent.

#### Studies related to obese human subjects of KOBS (human cohort 2)

As described previously (De Jesus et al. 2020), this study used data from 145 obese individuals (BMI ≥ 30) in the Kuopio Obesity Surgery (KOBS) Study (patient information is summarized in Supplementary table 4). PGE2 and PGF2a were measured by the Prostaglandin E2 Express ELISA Kit and Prostaglandin F2a ELISA Kit, respectively (Cayman Chemical, Ann Arbor, MI) according to the manufacturer’s instructions. For the correlation analysis, Box-Cox transformation on the values of metabolites (PGE2 and PGF2a) and clinical variables (BMI, triglycerides, fasting glucose, fasting insulin, HOMA-IR, and liver steatosis/NASH histology grades), which was appropriate for reducing skewness and to approximate normality (Box 1964) was performed. The bagged trees method was used for data imputation. Briefly, for each variable in the data, a bagged tree was created using all of the other variables. When a new sample has a missing variable value, the bagged model is used to predict the value. In theory, this is a powerful method of imputing.

#### Studies related to StopDia (human cohort 3)

Plasma samples were obtained from 73 individuals of the Stop Diabetes (StopDia) Study as described previously (Halali et al. 2022) (patient information is summarized in Supplementary table 5). Briefly, overnight (12 hours) fasting blood samples were obtainedand analyzed for glucose and insulin concentrations [fasting and 2 hours after the ingestion of 75 g glucose in an oral glucose tolerance test, glycated haemoglobin (HbA1c)] and fasting plasma total and lipoprotein lipid concentrations [total cholesterol, low-density lipoprotein (LDL) cholesterol, high-density lipoprotein (HDL) cholesterol and total triglycerides]. Matsuda index, as a measure of peripheral insulin sensitivity, was calculated using commonly used formulas. PGE2 and PGF2a were measured by the Prostaglandin E2 Express ELISA Kit and Prostaglandin F2a ELISA Kit, respectively (Cayman Chemical, Ann Arbor, MI) according to the manufacturer’s instructions.

#### Human brown adipocyte culture and differentiation

Immortalized human preadipocytes (Xue et al. 2015) were cultured in high-glucose DMEM containing 10% FBS. When cells reached confluence, brown adipocyte differentiation was induced by using induction medium (DMEM high-glucose medium with 10% FBS, 33 μM biotin, 17 μM pantothenate, 0.5 μM human insulin, 500 μM IBMX, 2 nM T3, 0.1 μM dexamethasone and 30 μM indomethacin) for 21 days. Once fully differentiated, cells were washed with PBS and RNA and protein were extracted for qRT-PCR and western blot analyses, respectively.

#### Human white adipocyte culture and differentiation

For differentiation of human white adipocytes, immortalized human preadipocytes (Xue et al. 2015) were induced using the same protocol and induction medium as described above for 14-18 days. Fully differentiated hWAT cells were co-cultured with freshly collected hBAT conditioned-medium, protein was extracted for western blot analysis.

#### Primary human myoblast culture and differentiation

Primary GIBCO® Human Skeletal Myoblasts (Thermo Fisher Scientific) were thawed and plated in DMEM supplemented with 2% horse serum. Cells were differentiated for 48 hours to form myotubes before being co-cultured with freshly collected hBAT conditioned-medium.

#### Primary human hepatocyte culture

Primary hepatocytes pooled from ‘5-Donor’ human hepatocytes (Thermo Fisher Scientific) were thawed and cultured in William’s E medium supplemented with primary hepatocyte supplements (Thermo Fisher Scientific) and HepExtend supplement (Thermo Fisher Scientific). Cell were washed with PBS and co-cultured with freshly collected hBAT conditioned-medium.

#### Cell line culture and differentiation

HepG2 human hepatocytes were purchased from ATCC and cultured with 5.5mM low glucose DMEM medium (Gibco) supplemented with 10% FBS, 1% penicillin, and 1% streptomycin in a humidified 5% CO_2_ incubator at 37 °C for 3 days to achieve 70% confluence before treatment.

C2C12 mouse skeletal myoblast cells were purchased from ATCC and cultured according to the manufacturer’s recommendations. When cells reached an 80-90% confluence they were induced to differentiate, by altering the growth medium to differentiation medium which consisted of DMEM medium, 5% HS (Horse Serum, Gibco, 16050114) and 1% P/S under the same culture environment.

#### Cell treatments

##### hBAT-conditioned media co-culture experiments

hBAT cells were differentiated for 21 days, washed with PBS and cultured in 1% BSA containing high-glucose DMEM medium for 24 hours. Fresh conditioned-media were collected for co-culture experiments. hBAT cells were stimulated with 100 nM of insulin for 15 min and cell pellets were collected for protein and RNA extraction (Figure 2G). Fresh conditioned-media were used for treating fully differentiated hWAT, primary human hepatocytes, and differentiated human myotubes for 16hr (Figure 2G). Recipients cells were washed with PBS and stimulated with 100 nM of insulin for 15 min and cell pellets were collected for protein extraction.

##### PG treatments

Overnight starved HepG2 cells (Figure S4A), differentiated C2C12 myotubes (Figure S4C), fully differentiated hWAT cells (Figure S4E), and fully differentiated hBAT cells (Figure S4G) were treated with 10, 100, 1000 nM of PGE2, PGD2, PGF2a, or 13,14-dh-15-keto-PGE2 overnight in the presence or absence of 500 nM of palmitate. Cells were washed with PBS and treated with 100nM of insulin for 15 min, cell pellets were collected for protein extraction.

##### PGE2 and PGF2a treatment with receptor antagonists

Overnight starved HepG2 cells, differentiated C2C12 myotubes, differentiated hWAT cells, and differentiated hBAT cells were pre-treated with 60 nM of ONO8711 (EP1 antagonist), 1600 nM of PF04418948 (EP2 antagonist), 80 nM of L826266 (EP3 antagonist), or 26 μm of AH23848 (EP4 antagonist) for 1hr, followed by 100 nM of PGE2 treatment overnight. In parallel, cells were pre-treated with 26 nM of OBE022 (FP antagonist) for 1hr before treatment with 10 nM of PGF2a. Cells were washed with PBS and stimulated with insulin (100 nM) for 15 min and collected for protein extraction. All the antagonists were purchased from Cayman Chemical.

##### Actinomycin D treatment

Differentiated sgNTC- and sgM14-hBAT cells were treated with 10 μg/μL Actinomycin D (Thermo Fisher) or DMSO for 0, 3, 6 or 12h. Differentiated Lenti-Control and M14OE-hBAT cells were treated with 10 μg/μL Actinomycin D (Thermo Fisher) or DMSO for 0, 3, 6 or 12h.

##### Cycloheximide treatment

Differentiated sgNTC- and sgM14-hBAT cells were treated with 100 ug/mL Cycloheximide (CHX) or DMSO for 0, 6 or 12 h, cell pellets were collected for protein extraction.

#### Transfections

##### Knock-out experiments

To generate stable M14 knockout in hBAT preadipocytes, three different Edit-R All-in-one Lentiviral human METTL14 single guide RNA (sgM14) and a Non-targeting control single guide RNA (sgNTC) was purchased from Horizon Discovery. After lentiviral transduction, hBAT preadipocytes were selected by puromycin treatment for 6 days to establish stable M14^KO^ colonies for adipocytes differentiation and functional experiments. Knockouts were validated by both qRT-PCR and western blot analyses.

##### Knock-down experiments

For transient knockdown of YTHDF1/2/3, reverse transfections were performed. Differentiated hBAT cells were mixed with Lipofectamine RNAiMAX Reagent (Life Technologies) and small interfering RNA complexes (Dharmacon) at a final concentration of 15 nmol/L siRNA according to manufacturer instructions. ON-TARGETplus Non-Targeting Control Pool D-001810-10-05, ON-TARGETplus Human YTHDF1 siRNA L-018095-02-0005, ON-TARGETplus Human YTHDF2 siRNA L-021009-02-0005, ON-TARGETplus Human YTHDF3 siRNA L-017080-01-0005 (Dharmacon, USA). hBAT cells were collected 96h post-transfection.

##### Overexpression experiments

For stable overexpression of METTL14 in hBAT preadipocytes, control and METTL14 overexpression lentiviral plasmids were designed. Overexpression was validated by both qRT-PCR and western blot analyses. Stable overexpression hBAT preadipocytes were expanded and differentiated for experiments including insulin stimulation, co-cultured experiments, and Actinomycin D treatment.

### METHOD DETAILS

#### Glucose and insulin tolerance tests

For glucose tolerance tests, mice were fasted overnight and injected via the intraperitoneal (I.P). route with glucose (2 g/kg body weight). Blood glucose levels were measured before (0 min), and 15, 30, 45, 60, 90 and 120 mins after injection. For insulin tolerance tests, mice were fasted for 6h, and injected I.P. with 1U/ kg insulin. Blood glucose levels were measured before (0 min), or 15, 30, 45, 60, and 90 mins after injection.

#### Food intake, energy expenditure and body composition measurements

For measuring daily food intake, mice were caged individually, the amount of feed was weighed and recorded before and after a 24hr interval. Daily food intake = the weight of feed provided – rest of the feed after 24 hr.

For indirect calorimetry, mice were individually housed in metabolic cages of a Comprehensive Lab Animal Monitoring System (CLAMS) at room temperature. After a 12h acclimation period, animals were exposure to acute cold exposure as described above in acute cold exposure section and then monitored for 72h to obtain energy expenditure measurements, the volume of oxygen consumption (VO_2_), and the volume of carbon dioxide production (VCO_2_). Body composition including total mass, fat mass, and lean mass was measured by Dual-energy X-ray absorptiometry (DEXA) scan, and lean mass was used for CLAMS outcome normalization.

#### RNA isolation and quantitative RT-PCR

High-quality total RNA (>200nt) was extracted using standard Trizol reagent (Invitrogen) according to manufacturer’s instructions and the resultant aqueous phase was mixed (1:1) with 70% RNA-free ethanol and added to Qiagen Rneasy mini kit columns (Qiagen) and the kit protocol was followed. RNA quality and quantity were analyzed using Nanodrop 1000 and used for reverse transcription using the high-capacity cDNA synthesis kit (Applied Biosciences). cDNA was analyzed using the ABI 7900HT system (Applied Biosciences) and gene expression was calculated using the ^ΔΔ^Ct method. Data was normalized to the expression of housekeeping genes. The sequences of primers used in this study are provided in Supplementary table 6.

#### Western blots and molecular analyses

Total proteins were harvested from tissue and cell lines lysates using M-PER protein extraction reagent (Thermo Fisher) and supplemented with proteinase and phosphatase inhibitors (Sigma), respectively, according to standard protocol. Protein concentrations were determined using the BCA standard protocol followed by the standard western immunoblotting protocol of proteins. The blots were developed using chemiluminescent substrate ECL (Thermo Fisher) and quantified using Image studio Lite Ver. 5.2 software (LICOR).

#### m^6^A immunoprecipitation and sequencing

To profile the m^6^A methylome, m^6^A sequencing (m^6^A MeRIP-seq) was utilized (X. Wang, Lu, et al. 2014). 1 μg of mRNAs were isolated from total RNA using a Dynabeads mRNA DIRECT purification kit (Thermo Fisher). Then, mRNA was adjusted to about 10 ng μl^−1^ in 100Lμl and fragmented using a Bioruptor ultrasonicator (Diagenode) with 30Ls on/off for 30 cycles. After that, 5 ul of each sample is saved as the ‘input’. m^6^A immunoprecipitation (m^6^A-IP) and library preparation were performed using the EpiMark N6-Methyladenosine enrichment kit (NEB). Input and RNA eluted from m^6^A-IP were used to generate the library using TruSeq stranded mRNA sample preparation kit (Illumina). All sequencing was performed on an Illumina HiSeq 6000 according to the manufacturer’s instructions.

#### Differential expression analysis for RNA-seq

As reported previously (De Jesus et al. 2019), the input library of m^6^A sequencing is essentially an mRNA sequencing library. Thus, we performed gene level differential expression analysis using the input libraries. Raw reads were trimmed using Cutadapt (4.2) and aligned to hg38 genome (GRCh38.107) and mm10 (GRCm38.102) using HISAT2 (2.2.1) (Kim, Langmead, and Salzberg 2015) and SAMtools (1.16.1). Aligned sequencing reads were analyzed using RADAR (0.2.4) (Zhang et al. 2019). Then we perform differential methylation analysis of count data using the R package DESeq2 (1.36.0) (Love, Huber, and Anders 2014). For both iBAT and hBAT samples, we used a cut-off P-value < 0.01 to select differential genes for pathway and gene ontology enrichment analyses. ConsensusPathDB45 and “Metascape” online tool were used to perform enrichment analysis.

#### Differential methylation analysis for m^6^A-seq

Using the R package RADAR (Zhang et al. 2019), approximately 25 million single-end 100-bp reads were generated for each sample. Counts were normalized for library size and IP counts were adjusted for expression level by the gene-level read counts of input libraries. Bins with average IP-adjusted counts lower than 10 in both control and M14^KO^ groups were removed. Then bins that were not enriched in IP were also filtered out. To construct PCA plots, we used the remove Batch Effect function in the limma package (Ritchie et al. 2015) to remove the batch, age, and body weight effects. To consider the pairing of m^6^A IP and input, we use the normalized, expression-level-(i.e. input)-adjusted, and low-read-count-filtered IP counts. Using Wald tests, we tested for significant effects of M14^KO^ on the m^6^A enrichment/depletion. We adjusted for multiple testing using the Benjamini-Hochberg false discovery rate (FDR) controlling procedure.

#### H&E staining and immunohistochemistry staining

For Hematoxylin and Eosin (H&E) staining, tissues of mice were fixed in 4% paraformaldehyde (PFA) overnight at 4°C, followed by dehydration in 70% ethanol. After dehydration, tissues were embedded in paraffin. Multiple 5 μm sections were prepared and stained with H&E following the standard protocol. Images were acquired using a Zeiss AxioImager M1 (Carl Zeiss). For immunostaining, paraffin-embedded tissues were deparaffinized twice in xylene and subsequently rehydrated. After heat-induced epitope retrieval using target retrieval solution (Dako), the tissues were blocked in PBS containing 10% goat serum with 0.1% Tween 20 for 60 min. After washing in PBS, slides from iBAT or iWAT tissues were incubated with various primary antibodies: rabbit anti-UCP1 (ab23841, Abcam, 1:250), antibody overnight at 4°C. The next day, slides were washed in PBS and incubated with goat anti-rabbit immunoglobulin G (IgG) (H + L) secondary antibody conjugated with Alexa Fluor 594 (Invitrogen, 1:200). Slides from eWAT were incubated with anti-F4/80 (ab300421, Abcam, 1:5000) for 30min at room temperature, followed by secondary antibody is LeicaDS9800 (Bond™ Polymer Refine Detection). Nuclei were stained using DAPI (4′,6-diamidino-2-phenylindole).

#### Measurement of adipocyte size

As reported previously (Kahraman et al. 2014), paraffin-embedded inguinal and epididymal white adipose tissue samples from control and M14^KO^ mice, cut into 5 μm sections, were stained with H&E. Digital images at 20 magnification were taken from non-overlapping fields, with 5 images per section, from a total of 6 animals per group. Images were analyzed using Cell Profiler software and the adipocyte pixel area was calculated to determine the adipocyte diameter. A total of 2,000 cells (CC group) or 1,250 cells (CL group) were analyzed per section.

#### Oil red O staining

Differentiated hBAT cells were washed twice with PBS and fixed with 10% buffered formalin for 30minutes at room temperature. Cells were then stained with a filtered Oil Red O solution (0.5% Oil Red O in isopropyl alcohol) for 2 hours at room temperature. Cells were washed several times with distilled water for final visualization.

#### Measurement of blood parameters

Insulin (Crystal Chem #90082), Adiponectin (Crystal Chem #80569), and Leptin (Crystal Chem # 90030) were measured by ELISA kits according to the manufacturer’s instructions.

#### PGE2 and PGF2a measurements

PGE2 and PGF2a concentrations were measured using the Prostaglandin E2 Express ELISA Kit and Prostaglandin F2a ELISA Kit, respectively (Cayman Chemical, Ann Arbor, MI) according to the manufacturer’s instructions.

#### *In vitro* glucose uptake assays

HepG2 cells, C2C12 cells, hWAT preadipocytes, hBAT preadipocytes were seeded in 96-well plate, cultured and differentiated as described above. Cells were pretreated with indicated antagonist for 1hr before treatment with 10 nM of PGF2a or 100 nM PGE2 overnight before glucose uptake was measured by Glucose Uptake-Glo™ Assay kit (Promega) according to the manufacturer’s instructions.

#### Untargeted LS-MS/MS signaling lipidomics

For lipidomic profiling, control and M14^KO^ mice were fasted overnight. Plasma, iBAT, iBAT conditioned-Krebs solution were collected (left panel in Figure 3A). Global lipidomics were also performed on cell pellets and cell-conditioned medium of sgNTC- and sgM14-hBAT cells (right panel in Figure 3A). The details of Lipidomic profiling were provided previously (Leiria et al. 2019; Lynes et al. 2017). For mouse samples, we performed moderated t-tests to discover lipids that were differentially abundant between M14^KO^ and control samples, with the adjustment of covariates age, body weight, and litters.

#### Quantification and statistical analysis

Data is displayed as means ± S.E.M. and P-values were calculated using either two-tailed Student’s t-test, or two-way ANOVA. All utilized statistical tests and ‘n’ numbers are presented in figure legends. For *in vitro* assays, ‘n’ corresponds to the number of experimental replicates. For animal assays or animal tissue extractions, ‘n’ corresponds to the number of mice used per genotype or condition. For human brown adipose tissue and plasma samples, ‘n’ corresponds to the number of donors used per group.

Figures have been generated using GraphPad Prism9, and BioRender.com.

## Supplemental Information titles and legends

**Figure S1, related to Figure 1. Ablation of Mettl14 in BAT corrects HFD-induced alterations in the major metabolic tissues**

(A) Protein level quantification of Mettl3, Mettl14, and Wtap in the iBAT of male control and *db/db* mice.

(B) Protein level quantification of Mettl3, Mettl14, and Wtap in the iBAT of male control and LIRKO mice.

(C) Western blot of Mettl14 expression in iBAT, eWAT, iWAT and from both female and male control and M14^KO^ mice.

(D) Western blot of Mettl14 expression liver and muscle from both female and male control and M14^KO^ mice.

(E) *Mettl14* gene expression in iBAT, iWAT, eWAT, liver, and muscle from both female and male control and M14^KO^ mice.

(F) Representative images of body size of control and M14^KO^ male mice fed with HFD.

(G) Representative images of gross appearance of control and M14^KO^ male mice fed with HFD.

(H-I) Representative images of H&E staining and IHC staining of UCP1 in iBAT (H) and iWAT (I) from control and M14^KO^ mice.

(J) Representative images of H&E staining and macrophage marker F4/80 IHC staining in eWAT from control and M14^KO^ mice.

(K) Representative H&E staining in liver and muscle from control and M14^KO^ mice.

(L and M) Adipocytes cell size distribution in iWAT (L) and eWAT (M) from control and M14^KO^ mice on LFD or HFD.

(N and O) Adiponectin and leptin levels in the serum from both female (N) and male (O) control and M14^KO^ mice.

(P) Western blot of Mettl14 and Ucp1 proteins in iBAT from LFD- and HFD-fed control and M14^KO^ male mice.

(Q) Rectal temperature of control and M14^KO^ male mice before and after cold exposure.

(R-T) Energy expenditure (Q), oxygen (O_2_) consumption (R), and carbon dioxide (CO_2_) production (S) of mice after acute (6-hr) cold (5 °C) challenge.

All samples in each panel are biologically independent. Data are expressed as means ± SEM. *p < 0.05, **p < 0.01, ***p < 0.001 by Two-tailed unpaired t-test (A, B, E, N, and O) and Two-way ANOVA (L, M, Q, R-T).

**Figure S2, related to Figure 2. Ablation of METTL14 promotes human brown adipocyte adipogenesis without impacting mitochondrial respiration and glycolytic function**

(A and B) The insulin-stimulated phosphorylation of Ir/Igf1rβ and Akt_S473_ in iBAT after injection of 1U insulin into the *vena cava*.

(C) METTL14 protein levels before and after differentiation in sgNTC-hBAT and 3 individual sgM14-hBAT cell colonies.

(D) Representative phase-contrast images (upper panels) and Oil Red O staining (lower panels) of differentiated sgNTC-hBAT and sgM14-hBAT cells.

All samples in each panel are biologically independent. Data are expressed as means ± SEM. *p < 0.05, **p < 0.01, ***p < 0.001 by Two-tailed unpaired t-test (B).

**Figure S3, related to Figure 3. Signaling lipids are responsible for the inter-organ communication induced by M14-deficient brown adipocytes**

(A) Representative western blot (right panels) and quantification (left panels) of pIR/IGF1Rβ and pAKT_S473_ in differentiated human white adipocytes treated with conditioned medium from sgNTC-hBAT or sgM14-hBAT, or denatured conditioned medium from sgNTC-hBAT (n=3 biological replicates). CM, conditioned medium.

(B-D) Principal component analyses (PCA) in untargeted lipidomics data of plasma (B), iBAT tissue (C), iBAT-conditioned Krebs solution (D) from control and M14^KO^ mice (samples are from 3 female and 6 male mice/group).

(E-F) Principal component analyses (PCA) in untargeted lipidomics data of hBAT cells (E), hBAT-conditioned media (F) from sgNTC and sgM14-hBAT (n=3 for each cell colony).

(G) Volcano plots of differentially down- (blue data points) and up-regulated (red data points) lipids in serum from M14^KO^ mice (samples are from 3 female and 6 male mice, log 2 (fold change threshold=2, *p* value threshold=0.05).

(H) Volcano plots of differentially down- (blue data points) and up-regulated (red data points) lipids in sgM14-hBAT cells (n=3 for each cell colony, log 2 (fold change) threshold=2, *p* value threshold=0.05).

(I) Venn diagram of all differentially abundant lipids in M14^KO^-iBAT, M14^KO^-iBAT conditioned Krebs solution, and M14^KO^-plasma (n=9 for each group).

(J) Venn diagram of all differentially abundant lipids in sgM14-hBAT conditioned medium and sgM14-hBAT (n=3-6 for each group).

(L) Venn diagram of all differentially abundant lipids in sgM14-hBAT conditioned medium, M14^KO^-iBAT conditioned Krebs solution, and M14^KO^-plasma (n=9 for mouse samples, n=3-6 for cell samples).

All samples in each panel are biologically independent. Data are expressed as means ± SEM. *p < 0.05, **p < 0.01, ***p < 0.001 by Two-tailed unpaired t-test (A).

**Figure S4, related to Figure 3. PGE2, PGD2, PGF2a, and 13,14-dihydro-15-keto-PGE2 are the M14^KO^-iBAT/hBAT secreted insulin sensitizers**

(A-D) Experimental scheme of culture, differentiation, and treatments in HepG2 (A), C2C12 (B), hWAT (C), and hBAT (D) cells. hWAT, human white adipocytes. hBAT, human brown adipocytes.

(E) Representative western blot (n=2 in total) of pIR/IGF1Rβ and pAKT_S473_ in HepG2, cells induced by PGE2 (10-1000 nM), PGD2 (10-1000 nM), PGD2 (10-1000 nM), 13,14-dihydro-15-keto-PGE2 (10-1000 nM), or insulin (100 nM) pre-treated without or with palmitate (500 nM).

(F) Representative western blot (n=2 in total) of pIR/IGF1Rβ and pAkt_S473_ in C2C12 cells induced by PGE2 (10-1000 nM), PGD2 (10-1000 nM), PGD2 (10-1000 nM), 13,14-dihydro-15-keto-PGE2 (10-1000 nM), or insulin (100 nM) pre-treated without or with palmitate (500 nM).

(G) Representative western blot (n=2 in total) of pIR/IGF1Rβ and pAKT_S473_ in hWAT, cells induced by PGE2 (10-1000nM), PGD2 (10-1000 nM), PGD2 (10-1000nM), 13,14-dihydro-15-keto-PGE2 (10-1000nM), or insulin (100 nM) pre-treated without or with palmitate (500 nM).

(H) Representative western blot (n=2 in total) of pIR/IGF1Rβ and pAKT_S473_ in hBAT cells induced by PGE2 (10-1000 nM), PGD2 (10-1000 nM), PGD2 (10-1000 nM), 13,14-dihydro-15-keto-PGE2 (10-1000 nM), or insulin (100 nM) pre-treated without or with palmitate (500 nM).

**Figure S5, related to Figure 3. Ablation of Mettl14 Improves Insulin Sensitivity Primarily through Prostaglandin E2 and Prostaglandin F2a**

(A, C, E, and G) Quantification of pIR/IGF1Rβ and pAKT_S473_ in HepG2 (A), C2C12 (C), hWAT (E), and hBAT (G) cells induced by PGE2 (100 nM) or insulin (100 nM) pre-treated with EP1/2/3/4 receptor antagonists ONO8711, PF04418948, L826266, and AH23848 (n=3 biological replicates).

(B, D, F, and H) Quantification of pIR/IGF1Rβ and pAKT_S473_ in HepG2 (B), C2C12 (D), hWAT (F), and hBAT (H) cells induced by PGF2a (10 nM) or insulin (100 nM) pre-treated with FP receptor Antagonist OBE022 (n=3 biological replicates).

(I) Quantification of SHIP1, SHIP2, PTEN, and PHLPP-1 in HepG2 cells pretreated with PGE2 in the presence of indicated EP1/2/3/4 receptor antagonists (n=3 biological replicates).

(J) Quantification of SHP1 and SHP2 in HepG2 cells pretreated with PGF2a in the presence of indicated FP receptor antagonist (n=3 biological replicates).

All samples in each panel are biologically independent. Data are expressed as means ± SEM. *p < 0.05, **p < 0.01, ***p < 0.001 by Two-tailed unpaired t-test.

**Figure S6, related to Figure 4. Administration of PGE2 and PGF2a improves insulin sensitivity independent of body weight in DIO mice**

(A and B) Locomotor activity (A) and ambulatory (B) activity of vehicle-, 25 mg/kg PGE2+PGF2a-injected HFD-fed mice after 3-week injection.

(C) Rectal temperature of vehicle-, 25 mg/kg PGE2+PGF2a-, or 50 mg/kg PGE2+PGF2a-injected LFD- or HFD-fed mice at room temperature.

(D) Average daily food intake of vehicle-, 25 mg/kg PGE2+PGF2a-, or 50 mg/kg PGE2+PGF2a-injected LFD- or HFD-fed mice after 3-week injection.

(E) Representative gross appearance of body size, adipose tissues, and liver in vehicle- or 25mg/kg PGE2+PGF2a-treated LFD-fed mice.

(F) Body weight of vehicle-, 25 mg/kg PGE2+PGF2a-, or 50 mg/kg PGE2+PGF2a-injected LFD- or HFD-fed mice at day 11 post injection where insulin tolerance test was performed.

(I-M) Quantification of pIr/Igf1rβ and pAkt_S473_ in liver (I), eWAT (J), iWAT (K), muscle (L), and iBAT (M) after 21-day injection of saline, 25 mg/kg, or 50 mg/kg PGE2+PGF2a, followed by 1U of insulin injection via *vena cava*.

(G) Rectal temperature of vehicle-, 25 mg/kg PGE2+PGF2a-, or 50 mg/kg PGE2+PGF2a-injected LFD- or HFD-fed mice after 6-hr cold exposure.

(H-J) Energy expenditure (H), oxygen consumption (I), and carbon dioxide production (J) of vehicle-, 25 mg/kg PGE2+PGF2a-injected LFD-fed mice after 6-hr cold exposure.

All samples in each panel are biologically independent. Data are expressed as means ± SEM. *p < 0.05, **p < 0.01, ***p < 0.001 by Two-tailed unpaired t-test (A-D, F, G) and Two-way ANOVA (H-J).

**Figure S7, related to Figure 5. Negative associations of circulating PGE2 and PGF2a with fasting plasma glucose, insulin, leptin, and triglycerides levels liver steatosis severity and in humans**

(A) Spearman correlation between the plasma levels of PGE2/PGD2 and fasting plasma glucose (FPG), fasting plasma insulin (FPI), leptin, and triglycerides in human cohort 1.

(B) Spearman correlation between the plasma levels of PGF2a and fasting plasma glucose (FPG), fasting plasma insulin (FPI), leptin, and triglycerides, in human cohort 1.

(C) Correlation between the plasma levels of PGE2 and glucose, insulin, and triglycerides in human cohort 2.

(D) Correlation between the plasma levels of PGF2a and glucose, insulin, and triglycerides in human cohort 2.

(E) Spearman correlation between the plasma levels of PGE2 and fasting plasma glucose (FPG), fasting plasma insulin (FPI), and triglycerides in human cohort 3.

(F) Spearman correlation between the plasma levels of PGF2a and fasting plasma glucose (FPG), fasting plasma insulin (FPI), and triglycerides in human cohort 3.

All samples in each panel are biologically independent. *p < 0.05, **p < 0.01, ***p < 0.001.

**Figure S8, related to Figure 6. M14 deficiency in iBAT or hBAT upregulates genes involved in the lipids metabolism pathways.**

(A) Top 20 enriched GOs and pathways of differentially upregulated genes in M14^KO^-iBAT versus control-iBAT.

(B) Top 20 enriched GOs and pathways of differentially downregulated genes in M14^KO^-iBAT versus control-iBAT.

(C) Enrichment of known m^6^A consensus motif RRACH in control- or M14^KO^-iBAT.

(D) Top 20 enriched GOs and pathways of intersected m^6^A-Hypo and upregulated genes in M14^KO^-iBAT versus control-iBAT.

(E) Differentially upregulated pathways in sgM14-hBAT versus sgNTC-hBAT.

(F) Heat maps of differentially upregulated genes involved in the prostaglandin synthesis and regulation pathway in M14^KO^-iBAT versus control-iBAT.

**Figure S9, related to Figure 7. METTL14-mediated m^6^A Installation Promotes Decay of PG synthases and their regulators mRNAs in a YTHDF2/3-dependent Manner**

(A) qRT-PCR analysis of *15-Pgdh* and *Ptgds* in the iBAT of male control and M14^KO^ mice. mRNA levels were normalized to β*-ACTIN* (n=8).

(B) Quantification of the indicated proteins in the iBAT of male control and M14^KO^ mice (B). GAPDH was used as a loading control (n=6).

(C) qRT-PCR analysis of *15-PGDH* and *PTGDS* in the sgNTC and sgM14 hBAT cells. mRNA levels were normalized to β*-ACTIN* (n=3).

(D) Quantification of the indicated proteins in the sgNTC and sgM14 hBAT cells. GAPDH was used as a loading control (n=3 for each cell colony).

(E) Quantification of indicated proteins in differentiated sgNTC or sgM14 hBAT cells incubated with 100 ug/mL Cycloheximide (CHX) for the indicated time. GAPDH was used as a loading control (n=3).

(F) Representative Oil Red O staining of Lenti-control and M14OE hBAT after 21-day differentiation at low (upper panels) and at high magnification (lower panels). n=3 for all groups. Scale bars, 100 μm.

All samples in each panel are biologically independent. Data are expressed as means ± SEM. *p < 0.05, **p < 0.01, ***p < 0.001 by Two-tailed unpaired t-test (A-D) and Two-way ANOVA (E).

