## Supplementary figures for "m^6^A mRNA Methylation in Brown Adipose Tissue Regulates Systemic Insulin Sensitivity via an Inter-Organ Prostaglandin Signaling Axis"

### Figure S1, related to Figure 1.

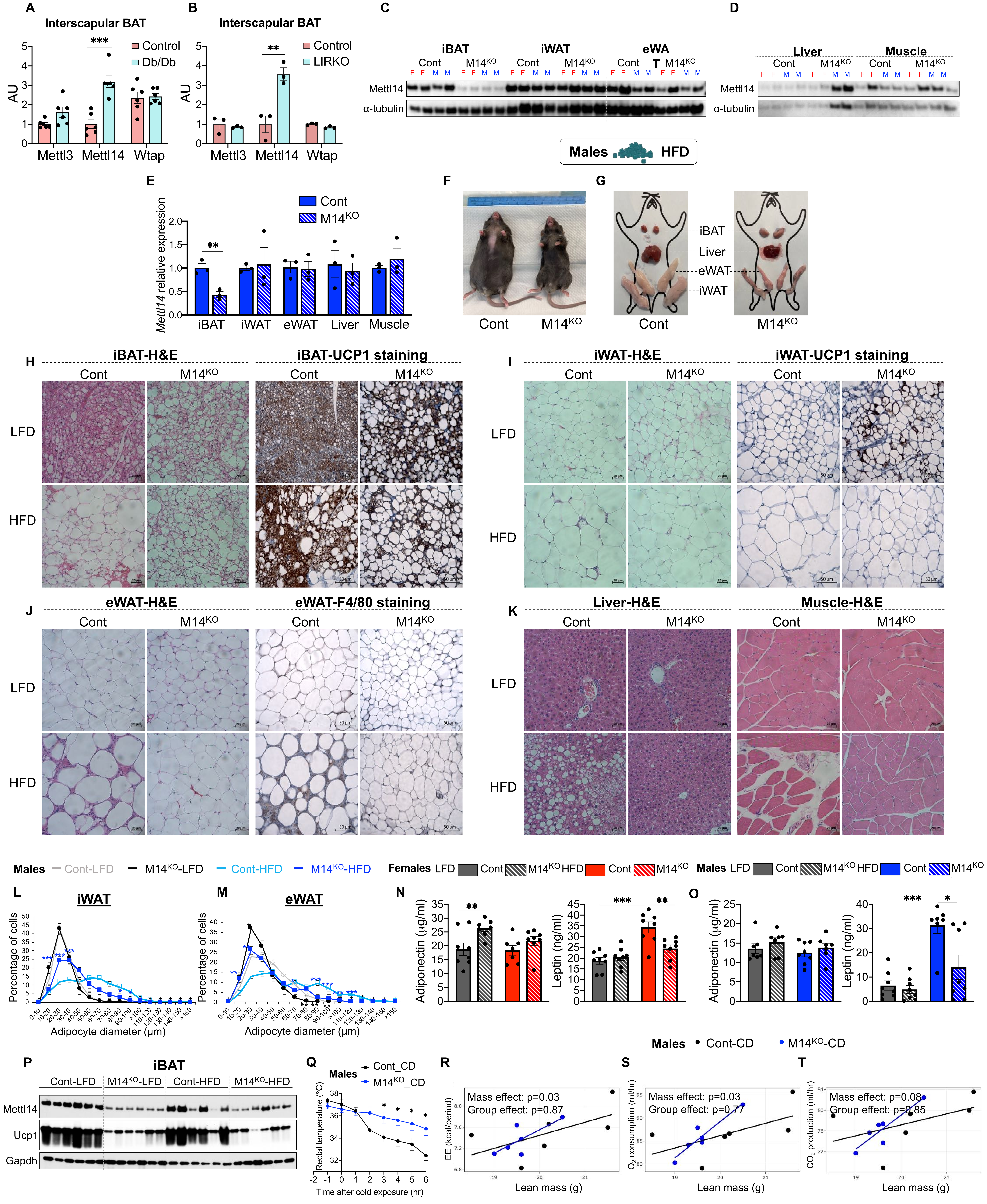

**Figure S2. Related to Figure 2.**

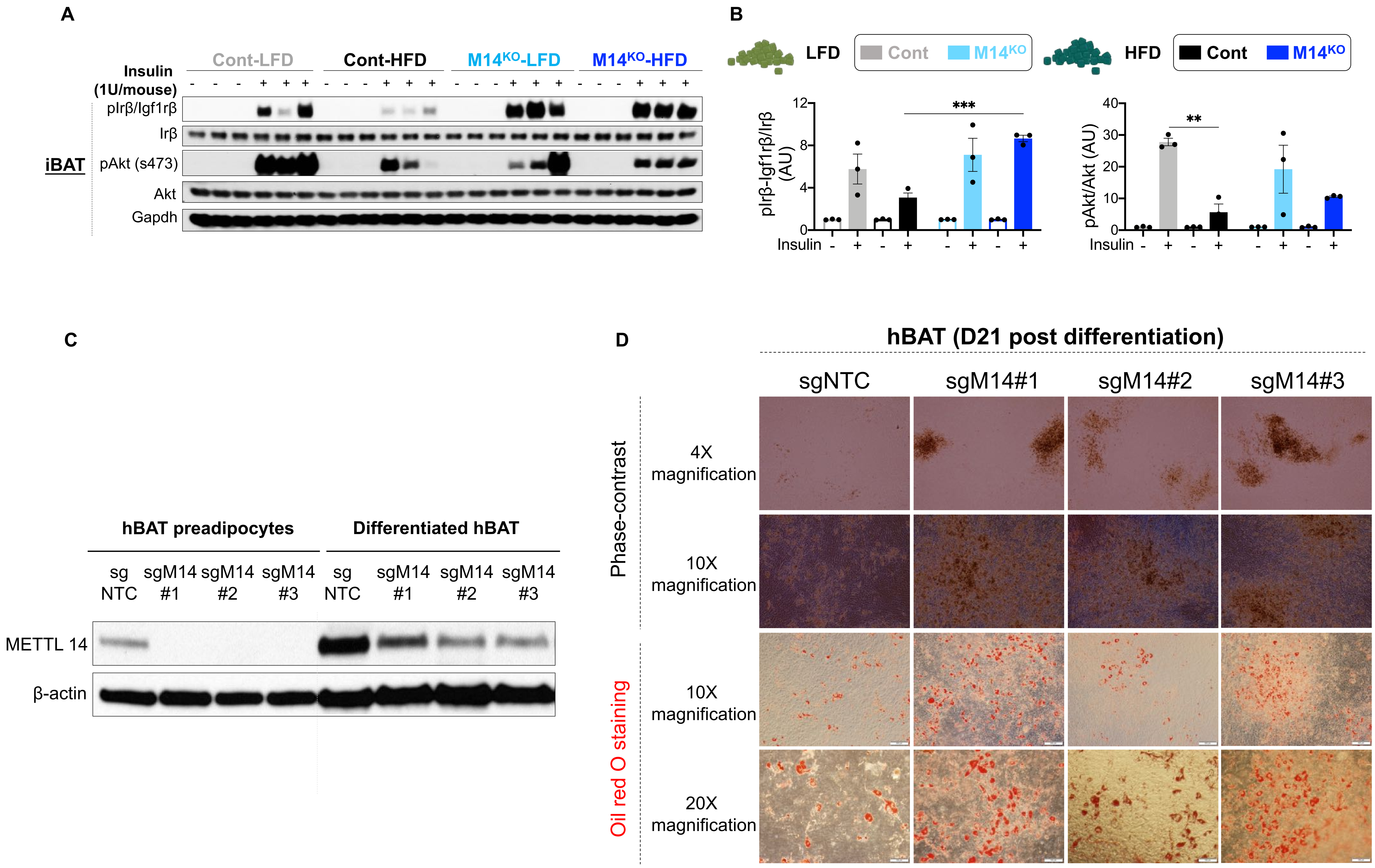

**Figure S3. Related to Figure 3.**

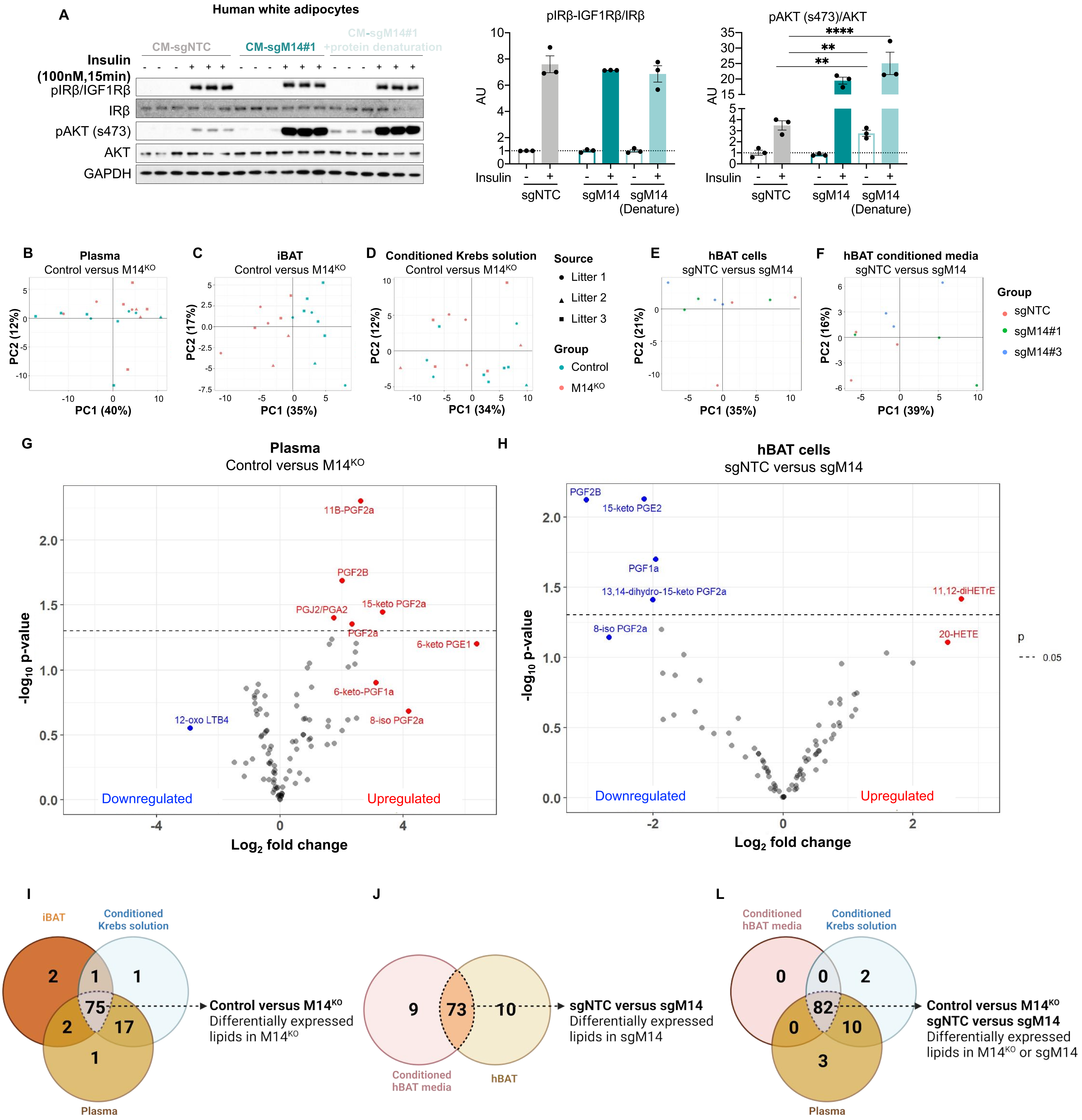

**Figure S4. Related to Figure 3.**

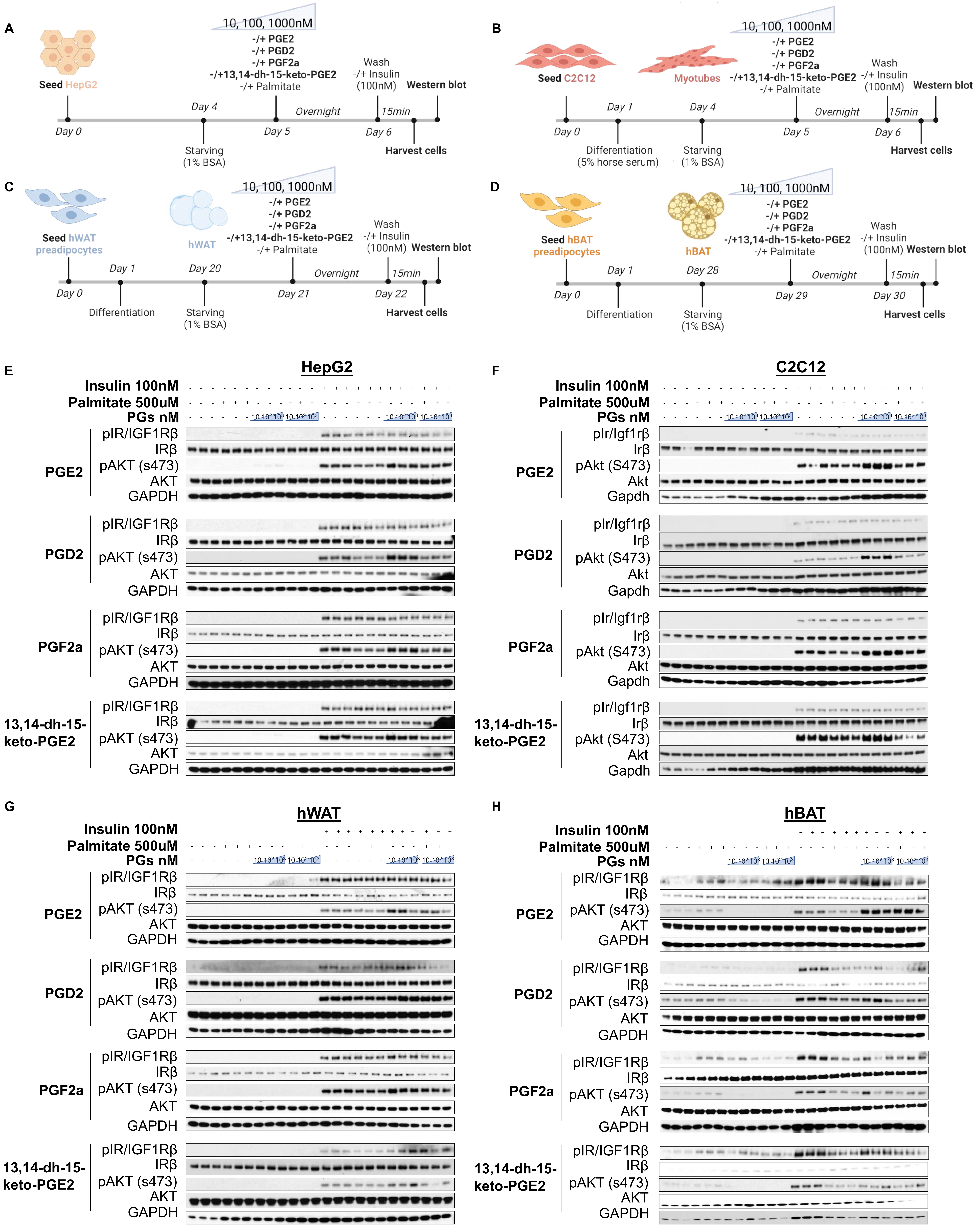

**Figure S5. Related to Figure 3.**

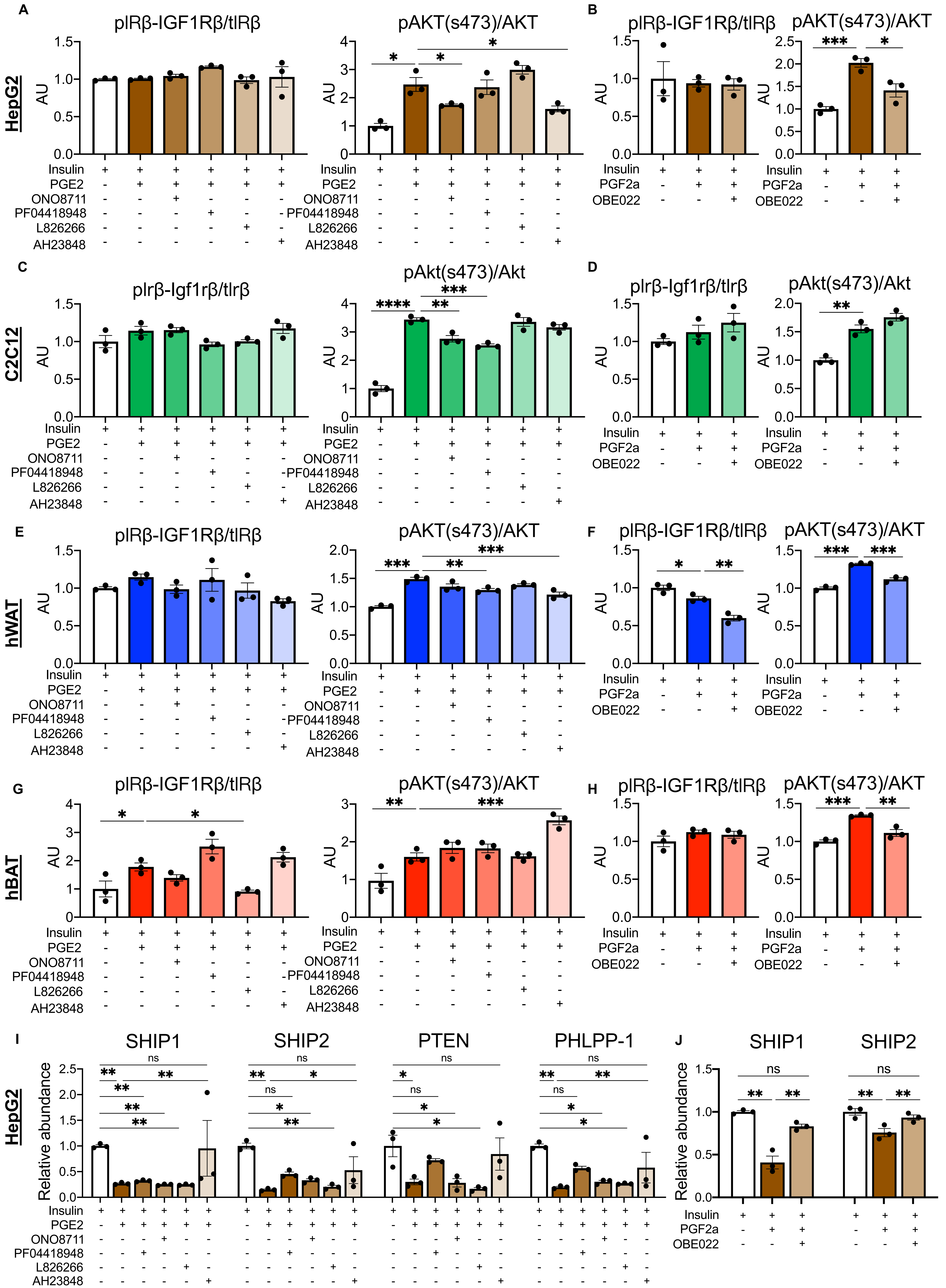

**Figure S6, Related to Figure 4.**

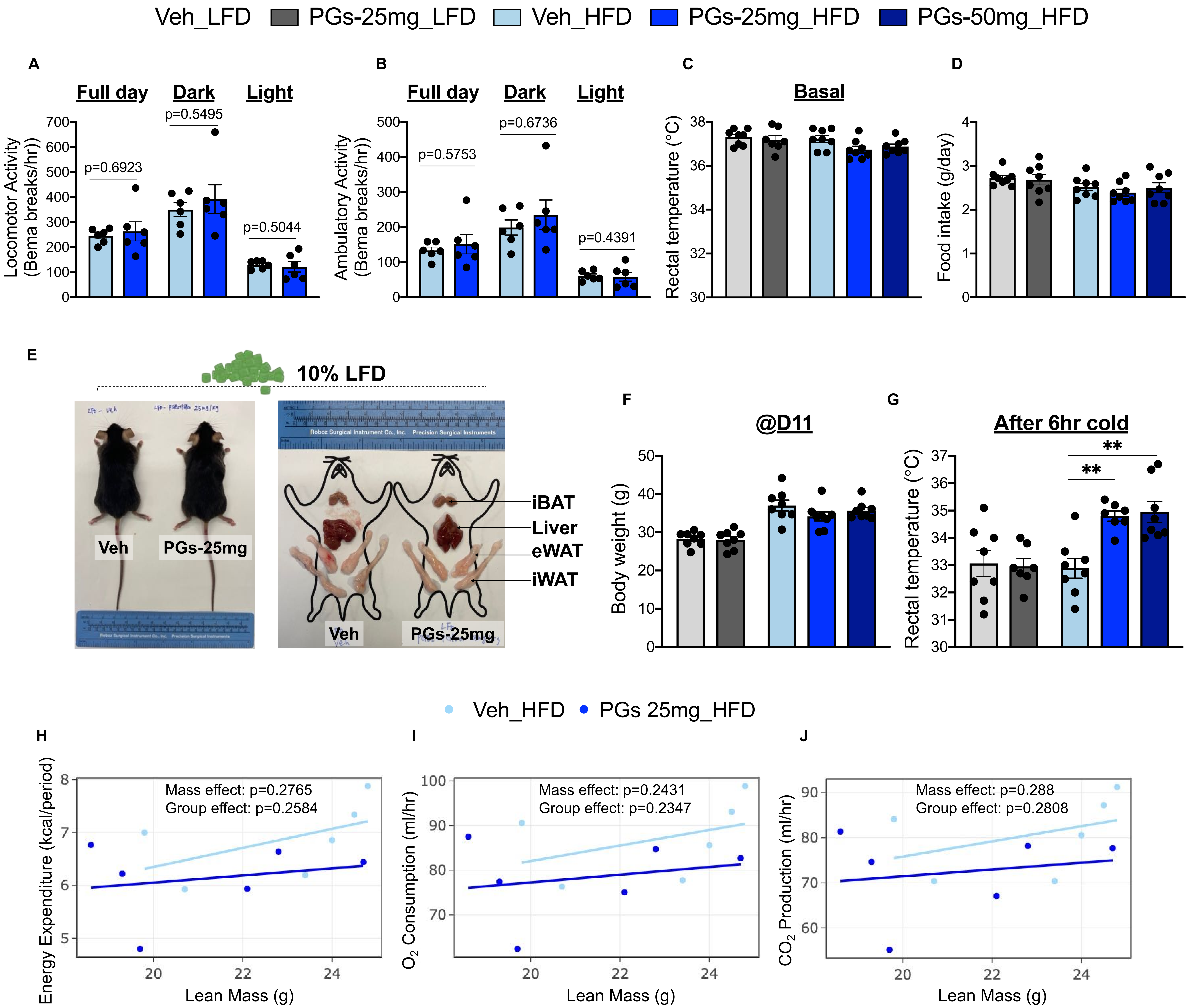

**Figure S7, Related to Figure 5.**

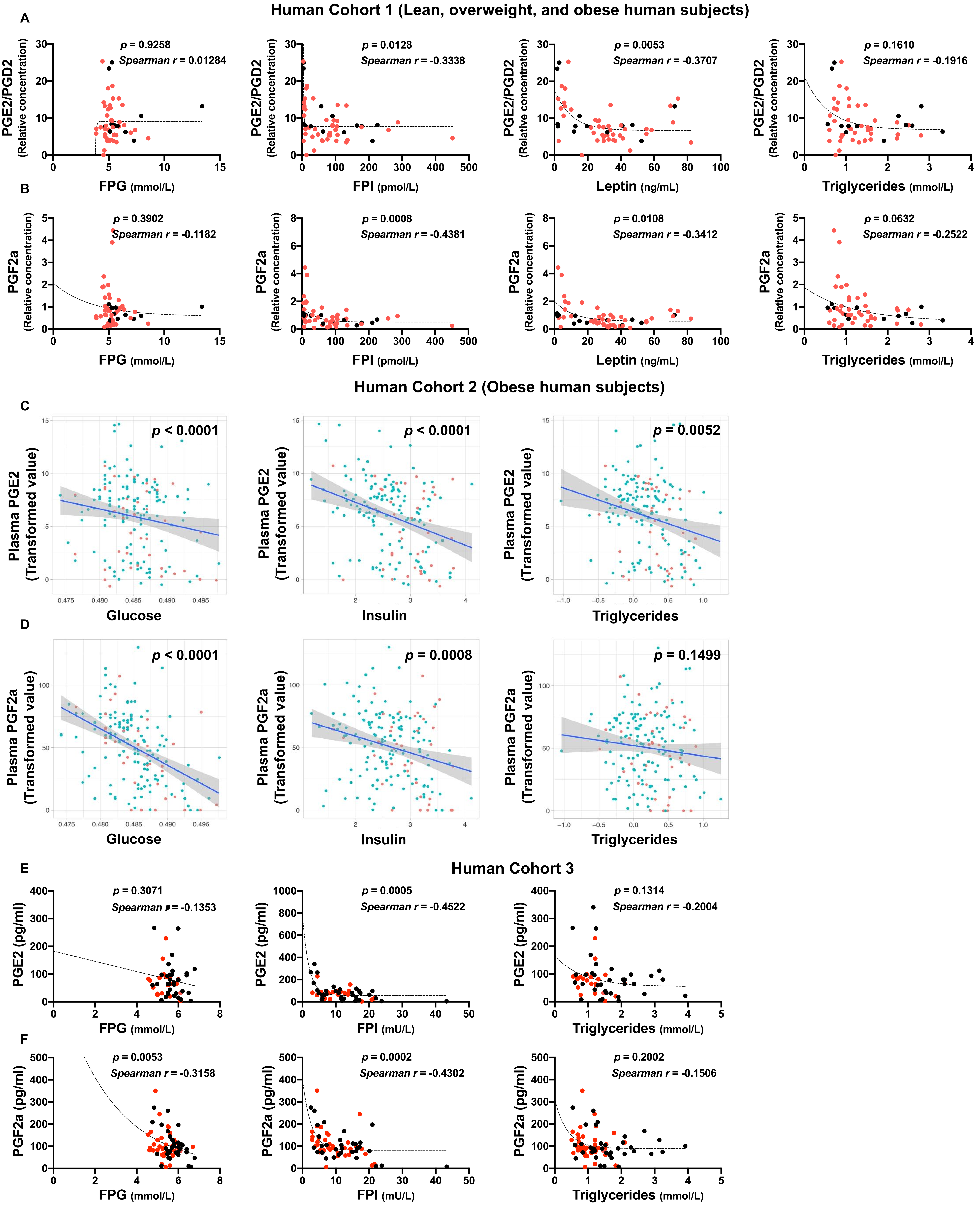

**Figure S8, Related to Figure 6.**

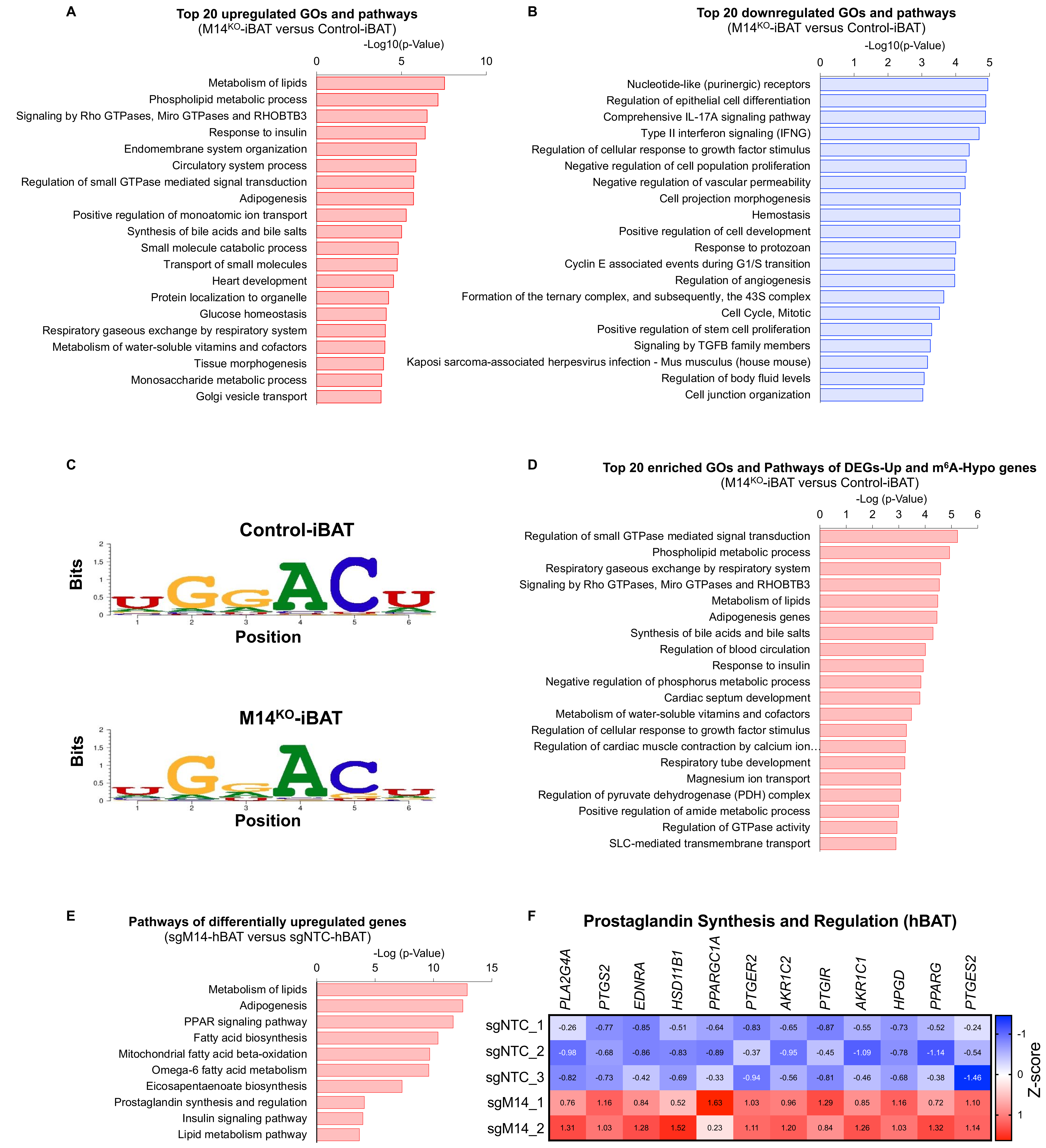

**Figure S9, related to Figure 7.**

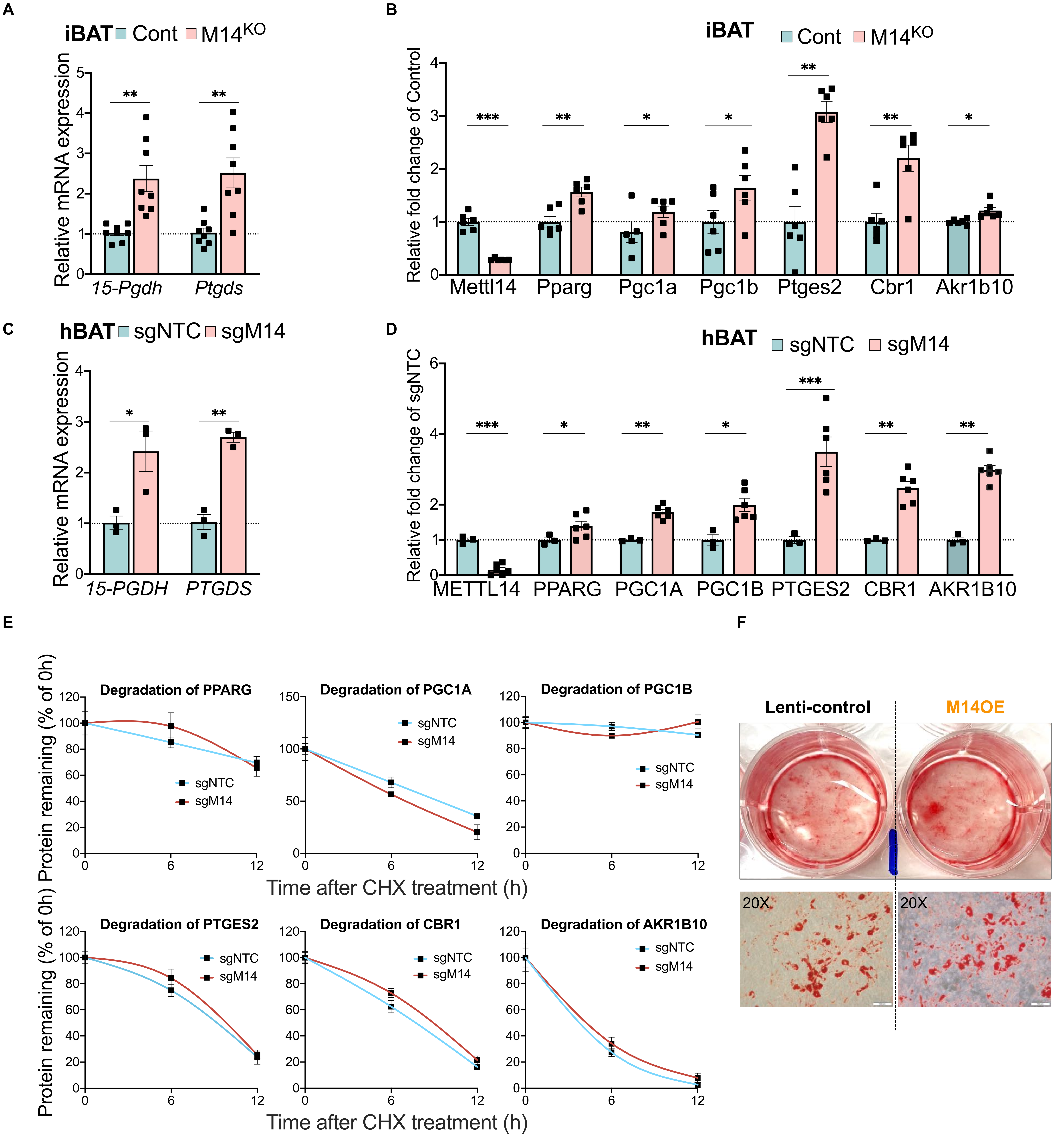

**Supplementary Table 1. related to Figure 1.** Clinical characteristics of The molecular, cellular, and genetic characterization of human adipose tissue and its role in metabolism human cohort.
.

| The molecular, cellular, and genetic characterization of human adipose tissue and its role in metabolism human cohort | n=13 |
|---|---|
| **Male/Female** | 6/7 |
| **BMI** | 20.1– 43.5 |
| **Age** | 21-74 |

**Supplementary Table 2. related to Figure 3.** $pAKT_{S473}$ increasing potency of candidate prostaglandins in HepG2, C2C12, hWAT, and hBAT. Notes: + indicates positive effects on $pAKT_{S473}$, − indicates no or negative effects on $pAKT_{S473}$.

| Cells / PGs | HepG2 | | C2C12 | | hWAT | | hBAT | |
|---|---|---|---|---|---|---|---|---|
| +insulin | insulin | insulin | insulin | insulin | insulin | insulin | insulin | insulin |
| +palmitate | | palmitate | | palmitate | | palmitate | | palmitate |
| PGE2 | + | + | + | + | + | + | + | + |
| PGD2 | + | + | + | + | − | + | − | − |
| PGF2a | + | + | + | + | + | + | + | − |
| 13,14-dh-15-keto-PGE2 | + | − | + | − | + | − | - | − |

**Supplementary Table 3.** related to Figure 5. Clinical characteristics of Human cohort 1 study.

| Human cohort 1 | n=55 |
|---|---|
| **Male/Female** | 13/42 |
| **BMI** | 17.5–75.4 |
| **Glucose (mmol/l)** | 3.9–13.4 |
| **Insulin (pmol/l)** | 3.8–451 |
| **HOMA-IR** | 0.1–25 |
| **Type 2 diabetes %** | 31% |
| **Normal glucose tolerance (NGT)** | 42% |

**Supplementary Table 4.** related to Figure 5. Clinical characteristics of the Kuopio Obesity Surgery (KOBS, Human cohort 2) study.

| Kuopio Obesity Surgery (KOBS) study (Human cohort 2) | n=145 |
|---|---|
| Male/Female | 38/108 |
| Age (years) | 48.06 ± 9.23 |
| BMI | 41.80 ± 4.59 |
| Glucose (mmol/l) | 6.07 ± 1.42 |
| Insulin (pmol/l) | 113.04 ± 65.73 |
| Cholesterol (mmol/l) | 4.29 ± 0.91 |
| HDL cholesterol (mmol/l) | 1.21 ± 0.31 |
| LDL cholesterol (mmol/l) | 2.47 ± 0.82 |
| Triglycerides (mmol/l) | 1.39 ± 0.69 |
| Type 2 diabetes (%) | 22.35 % |
| Histology (normal / simple steatosis / NASH) | 106/36/28 |

**Supplementary Table 5.** related to Figure 5. Clinical characteristics of the StopDia (Human cohort 3) study.
.

| (Human cohort 3) | n=145 |
|---|---|
| **Male/Female** | 39/39 |
| **Age (years)** | 26-60 |
| **BMI** | 20-35.6 |
| **Fasting Glucose (mmol/l)** | 4.56-6.79 |
| **Fasting Insulin (mU/l)** | 2.5-43.3 |
| **Cholesterol (mmol/l)** | 3.4-7.7 |
| **HDL cholesterol (mmol/l)** | 0.83-2.79 |
| **LDL cholesterol (mmol/l)** | 1.91-5.9 |
| **Triglycerides (mmol/l)** | 0.51-3.92 |
| **NGT/IFG/IGT** | 34/28/12 |

**Supplementary Table 6. List of primers and sequences. Related to Star methods.**

.

| Gene mane | Forward Primer | Reverse Primer |
|---|---|---|
| | **Human** | |
| **METTL3** | CTATCTCCTGGCACTCGCAAGA | GCTTGAACCGTGCAACCACATC |
| **METTL14** | CTGAAAGTGCCGACAGCATTGG | CTCTCCTTCATCCAGATACTTACG |
| **WTAP** | GTACAAGCTTTGGAGGGCAAGT | TGGACTTGCTTGAGGTACTGGA |
| **YTHDF2** | TAGCCAGCTACAAGCACACCAC | CAACCGTTGCTGCAGTCTGTGT |
| **YTHDF3** | GCTACTTTCAAGCATACCACCTC | ACAGGACATCTTCATACGGTTATTG |
| **PPARγ** | AGCCTGCGAAAGCCTTTTGGTG | GGCTTCACATTCAGCAAACCTGG |
| **PPARGC1α** | CCAAAGGATGCGCTCTCGTTCA | CGGTGTCTGTAGTGGCTTGACT |
| **PPARGC1β** | CGCTTTGAAGTGTTTGGTGAGATTG | GCTGGAAGGAGGGCTCGTTG |
| **PTGES2** | CCTCTATGAGGCTGCTGACAAG | ATCACACGCAGCACGCCATACA |
| **CBR1** | GCAAGTCAAAAGACATTGTTCTGG | TTGCCAAGGCACAAAGGACTGG |
| **AKR1B10** | GAGGACCTGTTCATCGTCAGCA | CGTCCAGATAGCTCAGCTTCAG |
| **15-PGDH** | TGGAGGTGAAGGCGGCATCATT | GAGCGTGTGAATCCAACTATGCC |
| **PTGDS** | AGCCCAACTTCCAGCAGGACAA | CAGACTTGCACATGGACAACGC |
| **β-ACTIN** | AGAGCTACGAGCTGCCTGAC | AGCACTGTGTTGGCGTACAG |
| | **Mouse** | |
| **Mettl3** | CAGTGCTACAGGATGACGGCTT | CCGTCCTAATGATGCGCTGCAG |
| **Mettl14** | AGAGTGCGGATAGCATTGGTGC | CTCCTTCATCCAGACACTTCCG |
| **Wtap** | AGTGCCTGGAAGTTTACGCCTG | GCTTCAAGCTGTGCAATACGGC |
| **Pparγ** | GTACTGTCGGTTTCAGAAGTGCC | ATCTCCGCCAACAGCTTCTCCT |
| **Ppargc1α** | GAATCAAGCCACTACAGACACCG | CATCCCTCTTGAGCCTTTCGTG |
| **Ppargc1β** | GGAGAAACCCCCTTTCCAGG | ACCTGAAGGTGCATCTGCTT |
| **Ptges2** | GGTAGACCTCTATGAAGCAGCC | CATCACTCGCAGCACACCATAC |
| **Cbr1** | CCTTCCACATTCAAGCAGAGGTG | CTGAGACTCACCATGCTGGACA |
| **Akr1b10** | GAGGACCTCTTCATCGTCAGCA | GCCAGTGGATTAGATACAGGTCC |
| **15-pgdh** | AAGCAAAACGGAGGTGAAGGCG | GAGCGTGTGAATCCGATGATGC |
| **Ptgds** | TCGCCTCCAACTCAAGCTGGTT | CCATGATCTTGGTCTCACACTGG |
| **β-actin** | CATTGCTGACAGGATGCAGAAGG | TGCTGGAAGGTGGACAGTGAGG |
